# Single-cell long-read mRNA isoform regulation is pervasive across mammalian brain regions, cell types, and development

**DOI:** 10.1101/2023.04.02.535281

**Authors:** Anoushka Joglekar, Wen Hu, Bei Zhang, Oleksandr Narykov, Mark Diekhans, Jennifer Balacco, Lishomwa C Ndhlovu, Teresa A Milner, Olivier Fedrigo, Erich D Jarvis, Gloria Sheynkman, Dmitry Korkin, M. Elizabeth Ross, Hagen U. Tilgner

## Abstract

RNA isoforms influence cell identity and function. Until recently, technological limitations prevented a genome-wide appraisal of isoform influence on cell identity in various parts of the brain. Using enhanced long-read single-cell isoform sequencing, we comprehensively analyze RNA isoforms in multiple mouse brain regions, cell subtypes, and developmental timepoints from postnatal day 14 (P14) to adult (P56). For 75% of genes, full-length isoform expression varies along one or more axes of phenotypic origin, underscoring the pervasiveness of isoform regulation across multiple scales. As expected, splicing varies strongly between cell types. However, certain gene classes including neurotransmitter release and reuptake as well as synapse turnover, harbor significant variability in the same cell type across anatomical regions, suggesting differences in network activity may influence cell-type identity. Glial brain-region specificity in isoform expression includes strong poly(A)-site regulation, whereas neurons have stronger TSS regulation. Furthermore, developmental patterns of cell-type specific splicing are especially pronounced in the murine adolescent transition from P21 to P28. The same cell type traced across development shows more isoform variability than across adult anatomical regions, indicating a coordinated modulation of functional programs dictating neural development. As most cell-type specific exons in P56 mouse hippocampus behave similarly in newly generated data from human hippocampi, these principles may be extrapolated to human brain. However, human brains have evolved additional cell-type specificity in splicing, suggesting gain-of-function isoforms. Taken together, we present a detailed single-cell atlas of full-length brain isoform regulation across development and anatomical regions, providing a previously unappreciated degree of isoform variability across multiple scales of the brain.

## Introduction

Studies characterizing the single-cell gene expression profiles of whole tissues and organisms have offered insight into the molecular makeup of complex tissues such as the brain^1–6^, evolutionary conservation of neuronal expression profiles^7–9^, and perturbations in neurological diseases^10, 11^. Early single-cell transcriptomic studies in the brain characterized the hippocampus and cortex, given the crucial roles of these brain structures in cognitive function. These seminal studies identified heterogeneous cell populations in mammalian neurodevelopment^1, 3, 4, 9, 12^. However, few brain single-cell studies consider mRNA isoforms. mRNA isoforms are strongly modulated in the mammalian brain, influencing processes such as cellular growth^13^, maturation^14–17^, migration^18, 19^, synapse formation^20, 21^, and activity patterns^22–26^. These properties of neuronal and non-neuronal cells are highly distinct between brain regions and change during development. Moreover, multiple diseases are associated with malfunctioned alternative splicing and may underlie regional vulnerabilities. While tissue-specific splicing is thought to have evolved across species^27–29^, there is an unmet need to identify conserved transcript elements defining the cellular diversity within highly specialized tissue such as the brain. Therefore, a comprehensive view of brain-region, cell-type, and developmental mapping of isoform usage at a single-cell resolution would aid our understanding of the brain in health and disease.

We and others have developed various single-cell short^30–32–^ and long-read^33–37^ technologies to study splicing. Long-read sequencing cDNAs end-to-end yields isoform profiles for thousands of single cells^34, 38–41^ thus allowing the quantification of isoforms within and between conditions. These studies in the mouse brain have shown that isoforms can define embryonic cell types^34^ and cell subtypes can differ in isoform expression and between two brain regions as early as postnatal day 7 (P7)^38^. However, the extent to which brain regions such as the thalamus, striatum, and cerebellum, which have distinct functions in motor control and coordination, differ in isoform expression for matched cell types is unknown. Furthermore, whether these regional differences differ in development or cell-subtype identity in defining splicing variation in neuronal and glial cell types is incompletely understood. Lastly, the degree to which any brain-region specific or cell-type specific isoform patterns are transient or maintained across development is an unsolved question.

Here, using an enhanced single-cell long-read method (ScISOr-Seq2), we investigate these questions in a comprehensive manner. We start by investigating full-length isoforms across three axes: multiple adult brain regions, cell subtypes, and developmental timepoints. We find that distinct types of neurons and glia show widespread and characteristic isoform variability along these three axes. Glia in thalamic and cerebellar regions exhibit especially strong transcription start site (TSS), PolyA site, and exon regulation. Distinct sets of exons display extremely high variability in their inclusion patterns across cell types, brain regions, and developmental time points, and cell-type specificity tends to be conserved in human brain. A strong splicing shift occurs in the hippocampal and cortical oligodendrocyte lineage after gene expression signatures already distinguish oligodendrocyte precursors from astrocytes. Fluctuations in splicing variation occur during mouse adolescence and exhibit a peak of neuronal subtype variability in the telencephalon, a period critical in the establishment and dissolution of splicing variability across all major cell types. These data showcase the importance of using long reads to capture a fuller picture of transcriptomic diversity in brain function.

## Results

### Short-read single-cell RNAseq identifies heterogenous cell populations

Based on our single-cell/nuclei isoform sequencing^33, 35^ studies, we devised ScISOr-Seq2 (Methods) to investigate brain-region specific, cell-type specific, and developmental-stage specific isoform regulation. Given the widespread transcription-mediated cell-identity establishment occurring in the telencephalon during postnatal development, particularly in the cortex and hippocampus which are crucial for memory and cognition, we first obtained single-cell 10x transcriptomics data from mouse hippocampus (HIPP) and visual cortex (VIS) at postnatal days 14, 21, 28, and 56 (n=16 samples: 2 replicates/age x 2 brain regions). To compare these substructures with diverse brain regions, we also obtained similar data from adult (i.e., P56) striatum (STRI), thalamus (THAL) and cerebellum (CEREB) (n=6 samples: 2 replicates/brain region x 3 brain regions). Filtering, quality control, short-read analysis^42, 43^ and integration-mediated batch-effect control^44^ yielded 204,725 cells (mean=9,300 cells/sample, Methods, **Table S1**). This allowed us to leverage the relatively high sequencing depth per single cell to identify brain cell types, namely neuronal, glial, vascular, and immune cells. Using marker genes and public databases^42, 43^, we defined three granularity levels for each cell: 1) Broad, e.g., neurons vs. glia, 2) medium cell type, e.g., excitatory vs. inhibitory neurons, and 3) cell subtype, e.g., layer2/3 vs. layer6 excitatory neurons (**Fig 1a**). Timepoint-specific UMAP embeddings described hippocampal neurogenesis with neuronal intermediate progenitors (NIPCs) and granule neuroblasts giving way to mature dentate gyrus granule neurons and pyramidal CA neurons. Oligodendrocyte progenitor cells (OPCs) and immature oligodendrocyte subtypes were enriched in pre-adulthood (**Fig 1b**). Similarly, mouse P56 brain-region specific UMAPs yielded broadly replicable cell types, albeit with certain brain-region specific neuronal populations such as layer-specific cortical excitatory cells, cerebellar granule cells, and type 1 vs. 2 medium spiny neurons (MSNs, **Fig 1c**). Gene expression of glial, immune, and vascular cells was more homogeneous across brain regions, e.g., cerebellar Bergmann glia aligned with telencephalic astrocytes in concordance with previous observations^45–47^ (**Fig 1 a,c**). Total cell numbers and cell subtypes were broadly consistent between samples and experiments, and microglia were often among the most abundant subtypes detected (**Fig 1d-e**). In total, we identified 34 cell subtypes and used these annotations to query differences in isoform expression between cell types arising from different brain regions and developmental timepoints with orthogonal long-read data.

**Figure 1.**
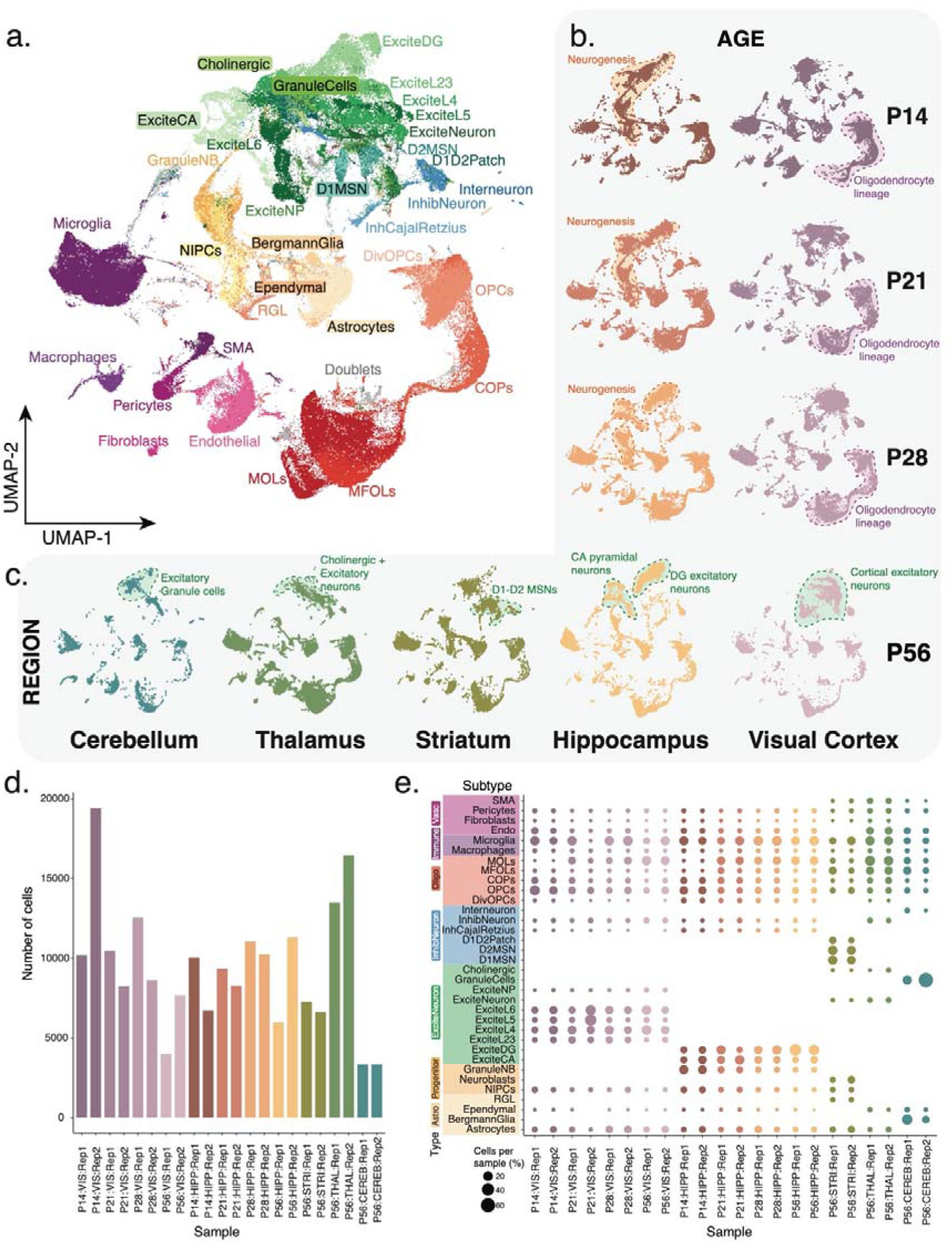
Summary of mouse brain cell subtype assignments by age and region. **(a)** UMAP embedding of all ∼200K cells. Each dot represents a cell which is colored according to its cell type of origin based on marker gene annotation. **(b)** Same UMAP representation from A but split by timepoint for the hippocampal (browns) and visual cortex (purples) lineage. **(c)** Same UMAP representation from A but split by the region of origin at P56. Blue: Cerebellum, Green: Thalamus, Olive: Striatum, Yellow: Hippocampus, Lilac: Visual Cortex. **(d)** Barplot depicting number of cells obtained from each single cell experiment. **(e)** Dotplot showing the percent of cells belonging to each cell subtype indicated on the Y-axis obtained from the samples on the x-axis. Color of the dots indicates sample of origin, size of dot indicates the percentage of cells belonging to a subtype per sample.

### Cell types are characterized by distinct regulation of isoforms across brain regions, development, and subtypes

Oxford Nanopore Technology (ONT) and PacBio HiFi sequencing yielded 250×10^6^ and 38×10^6^ barcoded long reads respectively for 395 cell clusters (e.g., P56:Thalamus:Replicate1:OPCs) obtained from the short-read analysis pipeline (Methods, **Table S2-3**). Using recent transcript-discovery software^48, 49^ high accuracy single-cell PacBio reads identified novel splice sites, enhancing the GENCODE annotation by 22.1% (40,184 transcripts). Over 67.3% of mapped, barcoded ONT reads (SD=4.51%, **Table S2**) represented multi-exonic transcripts with trustworthy splice sites, and ∼70% of these ONT transcript models corresponded to annotated or PacBio-derived transcripts (Methods, **Fig S1**). After removing UMI duplicates, all samples exhibited QC metrics commensurate with their sequencing depth, and these reads were used to calculate isoform abundance per cell cluster (**Fig S2**).

To parse the individual and/or coordinated roles of (i) developmental age, (ii) brain region, and (iii) cell subtypes in shaping the splicing programs for a given cell type, we determined the extent of isoform variability of the same cell type (progenitor, inhibitory neurons, etc.) across these three axes (Methods). Similar to TSS-exon-poly(A)-site contributions in the ENCODE project^50^, we represented the normalized age-subtype-region variability in a ternary plot. Each vertex of the triangle shows directed enrichment for the indicated axis of variability, while lines connecting the halfway points of the axes define a “center triangle” representing isoforms with broadly equal variability along all three axes (**Fig 2a**). Progenitor-cell isoforms varied more strongly by subtype than by age, suggesting that the switch from NIPCs to GranuleNBs is associated to strong isoform-mediated establishment of cell identity, regardless of developmental age. Perhaps unsurprisingly, as progenitors are less abundant in non-hippocampal adult regions, there is little variation in progenitor isoforms between brain regions (**Fig 2b**). Considering genes for which two isoforms were localized in distinct triangles in the ternary plots – indicating isoform regulation rather than gene regulation - we found enrichment for specific patterns. To isolate these patterns, we constructed a network diagram per cell type with nodes representing the axes, and the thickness of connecting lines indicating the number of such genes. One isoform largely exhibited progenitor-subtype or age variability while the other isoform(s) had uniform variability (center triangle) suggesting that different isoforms of a gene play distinct functional roles in establishing and maintaining progenitor identity (**Fig 2b, inset**).

**Figure 2.**
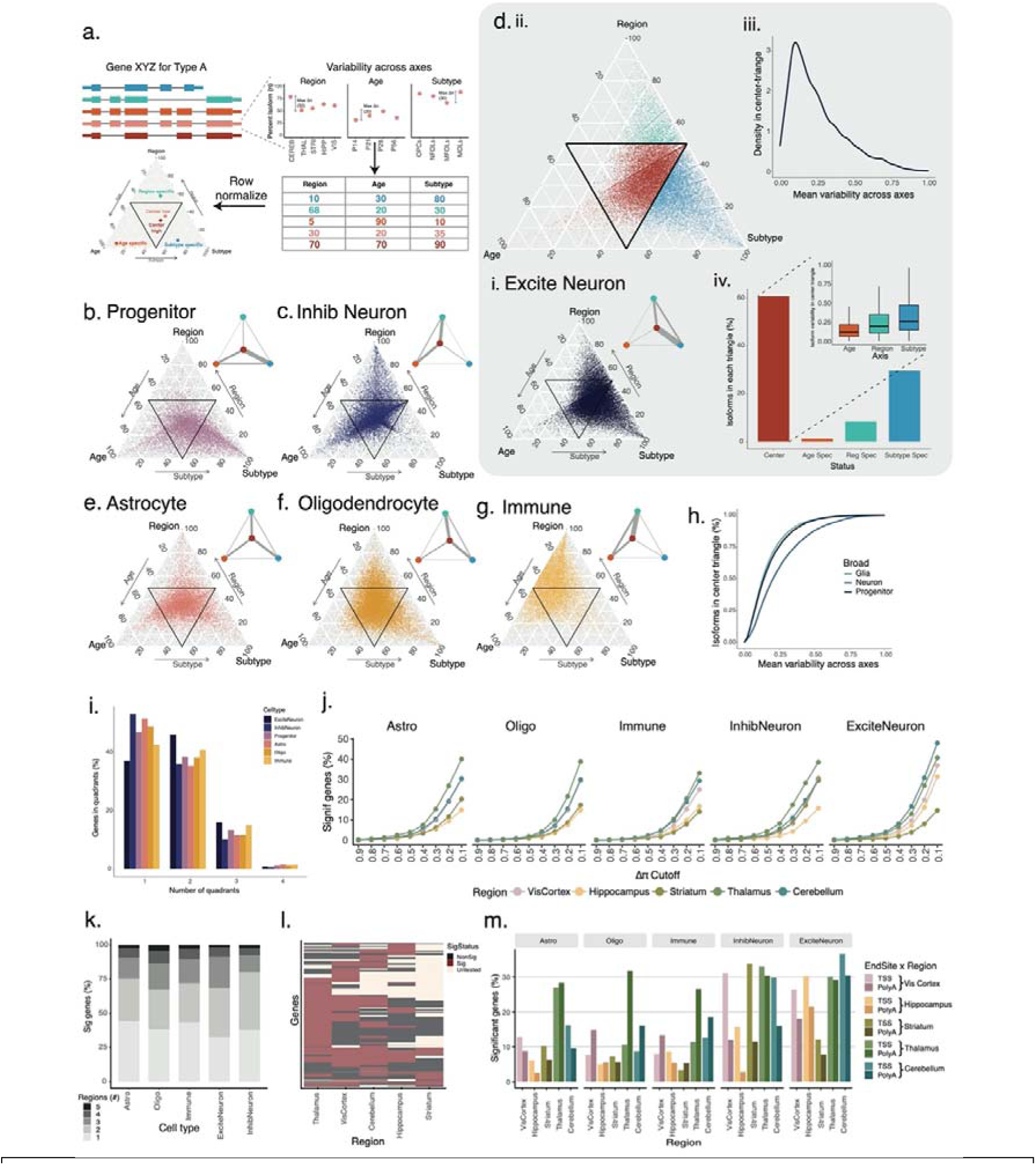
Distinct sources contribute to cell-type and brain-region specific isoform expression in the mouse brain. **(a)** Outline of full-length isoform variability across developmental age, brain region and cell subtypes. **(b)** Ternary plot of variability in three axes for progenitor cells. **(c)** Same as (b) for Inhibitory cells. **(d.i.)** Same as (b) for excitatory cells; (d.ii) points colored by origin of specificity: age-orange, region-teal, subtype-blue, center-red; (d.iii) Density of mean variability across three axes for each isoform; (d.iv) Fraction of isoforms in each triangle; inset shows variability of center-triangle isoforms **(e-g)** Same as (b) for astrocytes, oligodendrocytes & immune cells. **(h)** Comparison of mean variability for three broad cell types. **(i)** % of genes with isoforms in 1, 2, 3, or 4 triangles of variability. **(j)**. % of genes showing significant differences in isoform expression between astrocytes of one brain region vs. astrocytes of all other brain regions at 9 ΔΠ cutoffs (*left*). Neighboring four plots depict the same for oligodendrocytes, immune cells, inhibitory, and excitatory neurons. **(k)** % of genes with significant differential isoform expression (ΔΠ≥0.1) that are unique to one (light grey) or shared between multiple (darker grays) brain regions for each cell type. **(l)** Heatmap showing status (black–NonSig, Tan–Untested, Maroon–Significant) for each gene with significant differential isoform usage in one brain region compared to all others for immune cells. Foreground region indicated on the x-axis. **(m)** % of genes with significant differences in TSS/poly(A) site choice for astrocytes of a fixed brain region vs. astrocytes of all other brain regions at ΔΠ≥0.1 (left). Neighboring four plots depict the same for oligodendrocytes, immune, inhibitory, and excitatory cells.

Isoform variability was strong between ages and brain regions for inhibitory neurons – and in both cases stronger than for excitatory neurons. Additionally, increased abundance of excitatory neurons allowed the measurement of subtype-specific variability within one region, which was possible to a lesser extent for inhibitory subtypes (**Fig 2c, 2d.i.**). Indeed, for excitatory neurons, strong variation across subtypes for one isoform was frequently accompanied by uniform variation across all three axes for the other isoform (Network diagram **Fig 2d.i.**). Due to the normalized matrix of variability being represented, center-triangle isoforms by definition have similar variability across all three axes – with variabilities being either all high, or all low (ref Fig 2a, **Fig 2d.ii.**). For excitatory neurons, most such center-triangle isoforms had consistent low variability, however a few showed consistent high variability (**Figure 2d.iii.**). Additionally, despite their similar variability across three axes, excitatory neuron isoforms in the center triangle tend to display subtype specificity, mirroring the trend in the entire excitatory neuronal population (**Fig 2d.iv.**). A similar observation was made across all cell types (**Fig S3**). Among glia, both astrocytes and oligodendrocytes showed complex variability patterns along regions, ages and subtypes, but oligodendrocytes had stronger variability across regions and subtypes (**Fig 2e-f**). The microglial subtype dominated the immune population, where little variability between subtypes was observed. However, strong regional isoform variability characterized immune cells. (**Fig 2g**). In summary, distinct cell types exhibit distinct preferences for isoform variability across brain regions, age, and cell subtypes pointing to a complex interplay of the three axes of phenotypic definition.

Considering overall patterns, isoforms had higher mean variability across the three axes in neurons than in glia or progenitors, pointing to more complex neuronal regulation (**Fig 2h**). Approximately 75.4% of genes had isoforms in distinct triangles and thus showed isoform variability independent of gene variability along at least one axis for one or more cell types. Moreover, 53.26% of genes showed isoforms in three triangles, revealing isoform variability along two or three axes. However, this analysis does not allow for the comparison of isoform patterns of individual genes between different medium-level cell types (excitatory/inhibitory neurons, astrocytes, etc). We found that at least 36.6% of genes showed a variability in isoform usage between these cell types in addition to regulation along another axis (Methods, **Fig S4a**). Within a fixed cell type, between 43.4% (in inhibitory neurons) and 53.34% (in excitatory neurons) of genes had isoforms in distinct triangles. Hence, excitatory neurons display isoform regulation not recapitulated in other cell types (**Fig 2i**). In addition to a third of genes showing multi-axial variability in most cell types, ∼66% genes showed this hyper variability only in restricted cell types (**Fig S4b**). The gene *Rufy3*, which plays a role in neuronal polarity^51^, axon growth^52^, and synaptic plasticity^53^, exemplifies cell-type specific isoform variability along multiple axes. Three of the six annotated transcripts exhibit developmental variability in distinct cell types (progenitors, inhibitory neurons, and immune cells), two isoforms exhibit subtype and regional variability in astrocytes, while one isoform (Rufy3-207) is consistently found in the center-triangle (**Fig S5**). This points to a previously underappreciated complexity of isoform expression for genes such as *Rufy3* in conferring specialization across neurodevelopment, cell (sub)types, and brain regions.

Given the widespread brain-region variability in isoform expression within individual cell types, we systematically tested genes for altered isoform expression for matched cell types comparing one brain region to all other brain regions at P56 using our isoform tests and differential isoform quantification^53^ (ΔΠ, Methods). Thalamic and cerebellar astroglia showed strong specialized isoform expression compared to other brain structures. This is particularly interesting since cerebellum contains specialized Bergmann glia that are both morphologically and functionally distinctive^54^. This high splicing specialization supports alternative splicing as an important influence on brain anatomical region morphology and function. At modest differences of 10% isoform usage between conditions (ΔΠ=0.1, FDR ≤ 0.05, Methods), ∼40% of tested genes show a significant difference in isoform abundance between thalamic astrocytes and all other sequenced astrocytes. Broadly similar observations arose for oligodendrocytes and immune cells. Neurons had more differentially expressed isoforms, with medium spiney neurons (MSNs) contributing to high brain-region specificity of striatum inhibitory neurons, and distinct hippocampal pyramidal cells contributing to region specificity in excitatory neurons. These observations remained true, albeit lower, for increased ΔΠ values across all cell types. However, for ΔΠ≥0.5 few brain-region differences arose indicating that regional differences in isoform expression arise from smaller modulations across many genes (**Fig 2j**).

While a large percentage of genes exhibited unique regional signatures, most genes showed distinct isoform expression in two or three regions, and rarely in four or five (**Fig 2k**). Consistent with the observation in Fig. 2g, immune populations had high regional specificity, and thalamus was an outlier in differential isoform expression, in which a high number of significantly differentially spliced genes were either non-significant or untestable in other brain regions (**Fig 2l, Fig S6**). Region-specific isoform expression of immune and oligodendrocyte populations had more changes in Poly(A)-site as opposed to TSS usage. Compared to glia, neurons exhibited higher levels of differential TSS and PolyA site usage between regions, and neuronal TSS usage was more region-specific than that of glial or immune types (**Fig 2m**). In summary, in addition to isoform regulation across development and between distinct cell (sub)types within an anatomical structure, the same cell type present in different brain regions leverages distinct splicing patterns, alternative TSS and PolyA sites, pointing to brain-region specificity of isoforms tied to distinct structure, localization, and function.

### Marker exons delineate cell-type and brain-region specific splicing patterns conserved in human

To define precise transcript elements underlying brain-region and developmental splicing programs in the mouse brain, we focused on individual exons as opposed to full-length isoforms. We considered exons alternatively included in at least one brain region or time point and calculated the percent-spliced-in (Ψ) values for four main cell types (astrocytes, oligodendrocytes, excitatory neurons, inhibitory neurons) after averaging both replicates (Methods). Similar to the three axes of isoform variability, an exon’s Ψ value can vary along the triad of (a) cell subtypes, (b) matched cell types at different ages, or (c) brain regions (P21:Hippocampus:Oligo, n=44 triads, **Fig S7**). Pairwise correlations of Ψ values of such triads separated neuronal from non-neuronal populations, and all adult (P56) astrocytes clustered together regardless of region of origin. However hippocampal excitatory neurons clustered together regardless of age. Thus, unifying programs of age and/or brain region do not dictate splicing of distinct cell types (**Fig S8**).

We next compared exon inclusion between triads to isolate splicing programs. This yielded 4557 exons where a 25% difference in exon inclusion (ΔΨ≥0.25) was observed in at least one comparison, which we termed **h**ighly **v**ariable **ex**ons (hVEx, Methods). Hierarchical clustering showed that the highest ΔΨs corresponded to neuron vs. non-neuron comparisons – corresponding to a vertical split in the heatmap (*top left*, bipartite network diagram). On average, moderate ΔΨs corresponded to comparisons between two neuronal or two glial triads (*top right*, fully-connected network diagram). Many comparisons in the right half of the heatmap corresponded to differences between finer cell subtypes, or to brain-region and developmental differences of a matched cell type as indicated by self-loops in the network diagram. Clustering along rows defined four exon groups (A1-A4, **Fig 3a**), whose genes harbor largely non-overlapping functional ontologies: A1 and A4 hVEx have high ΔΨs for a few comparisons and low ΔΨs for most others. These exons’ genes are linked to regulatory roles for transcription, histone methylation and acetylation, and synaptic signalling. Conversely, hVEX exhibiting high ΔΨs in many comparisons, especially in the left half of the heatmap (A2, A3) belong to genes implicated in protein localization and neurotransmitter transport to the synapse. Together with the clean neuron versus glial split in the left half of the heatmap, these observations emphasize the role of synaptic isoforms rather than simple changes in expression of synaptic genes in establishing neuronal and glial identities (**Fig S9**).

**Figure 3.**
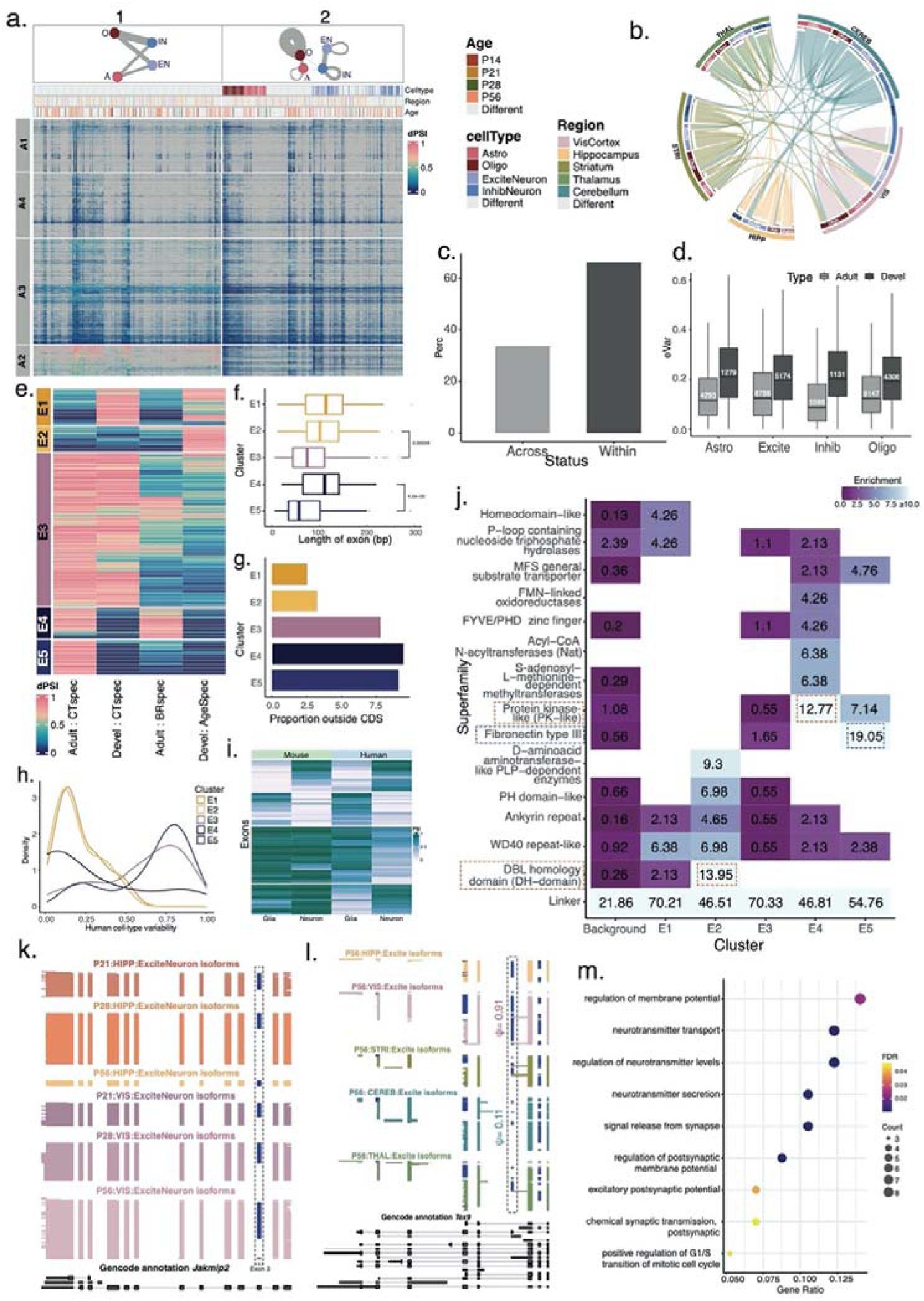
Marker exons underlying distinct splicing programs correlate with function and are conserved in human. **(a)** ΔΨ heatmap for pairwise cluster comparisons (columns) and exons (rows) where ΔΨ≥0.25 in at least one comparison. **(b)** Circos plot for connections between triad pairs (P56). Outer concentric ring: brain region; inner concentric ring: cell types. At constant age (P56), triad is defined by brain region and cell type. Triads connections indicate that HVExs are present in this comparison. Connection thickness indicates number of HVExs detected in comparison. Connection color indicates brain region of origin but cross-brain region-comparison colors are random. **(c)** % of HVEXs whose variability stems from a comparison of a matched cell type across brain regions or from two cell types in the same region **(d)** Maximal ΔΨs for matched cell types across brain regions and across developmental timepoints **(e)** Heatmap of EVExs (rows) and axes of variations (columns: adult cell-type specificity, developmental cell-type specificity, adult brain-region specificity and developmental specificity of a matched cell type) Five EVEx classes indicated in bar on the left **(f)** Length distribution of 5 EVEx classes **(g)** Non-coding fraction for 5 EVEx classes **(h)** Cell-type variability of mouse EVExs in human hippocampus **(i)** Heatmap of neuron and glia Ψ values for mouse (left) and human (right) for exons that have high cell-type specificity in human **(j)** Protein domains enriched in 5 EVEx classes **(k)** Cluster-resolved single-cell long reads for *Jakmip2* gene. Each line is a single cDNA molecule. Blue exons: alternative exons. Top 3 tracks: Hippocampal excitatory CA isoforms for P21, P28 & P56. Next 3 tracks: Visual-cortex excitatory isoforms from P21, P28 & P56. Bottom black track: Gencode annotation **(l)** Similar as (j) for *Tex9* gene with tracks colored for brain region of origin for P56 excitatory neurons. (k-l) Plotted with ScisorWiz **(m)** GO biological process annotations for EVEx in E4 from (e)

By definition, each hVEx arises from one or more triad comparisons with highly different Ψ values. Considering the triads giving rise to hVEx, we found that while most came from cluster comparisons within one brain region, a cell type could also show variability across brain regions (**Fig 3b**). Indeed, 33.71% of these comparisons corresponded to a matched cell type between two brain regions, and 66.29% to a comparison within the same brain region (**Fig 3c**). We then compared brain-region and developmental Ψ variability for matched cell types. Developmental variability exceeded variability between adult brain regions (median ΔΨ=0.115 and 0.197 resp., Wilcoxon rank-sum-test p<2.2e^-16^). This was equally true for each major cell type, suggesting that cell types extensively modulate exon inclusion over time but reach homeostasis in adulthood for many genes (**Fig 3d**).

To delineate the markers of this extensive modulation in matched cell types across brain regions and development and between cell types at a given developmental timepoint or brain region, we defined **e**xtremely **v**ariable **ex**ons (EVEx, ΔΨ≥0.75 between two triads, Methods) as representing potential candidates for functionality. Specifically, we determined five exon groups (E1-E5): 89 exons in E1 with cell-type specific inclusion during development but not in adulthood; 60 exons in E2 with temporal variability. The largest group, E3, had 373 exons with cell-type specificity in development and in adulthood. E4 included 75 exons with brain-region variability between matched cell types, combined with cell-type specificity within a brain region. Thus, variability between matched cell types of different brain regions largely implied cell-type specialization within one region. Lastly, E5 contained 84 exons with cell-type specificity acquired only in adulthood (**Fig 3e**). Exons in E3 and E5, i.e., those with adult cell-type specificity, regardless of earlier cell-type specificity, were markedly shorter, suggesting a link to microexons^35, 55^. Exons in E1, E2 (transient cell-type specificity in development) and E4 (brain-region specific regulation) were all longer (**Fig 3f**, two-sided Wilcoxon rank sum test p=1.62e^-10^). Moreover, exons with adult cell-type specificity (E3, E4, E5) harbored more non-coding sequence (**Fig 3g**, Fisher’s exact test p=1.5e^-4^).

To evaluate the transferability of EVEx to human tissue, we obtained single-cell long-read data for six adult human hippocampi and compared cell-type specific exon inclusion between human and mouse (Methods, **Figure S10**). Among the EVEx, 24.09% (seen in E5) to 48.31% (in E1) of exon sequence and boundaries were conserved between species and were sufficiently expressed in our human hippocampal data (Methods, **Fig S11a**). Among those, exons with cell-type specificity in mouse (E3, E5) tended to also exhibit high cell-type specificity in human (**Fig 3h**, Wilcoxon p<2.2e^-16^). Probing the transferability from human to mouse revealed that among exons that were highly cell-type specific in human hippocampus, ∼42.6% showed similar cell-type specificity in mouse hippocampus. However, ∼57.4% of human cell-type specific and almost 82.7% of invariable alternative exons in human were constitutively included in mouse tissue, suggesting that human brains evolved some gain-of-function exons and the expression patterns of these cannot be modelled well in mouse (Methods, **Fig 3i**, **Fig S11b**).

We next sought to estimate the functional impact of EVEx inclusion patterns by determining affected protein domains (Methods). The “inter-domain linker” was most frequent in all EVEx groups. Inter-domain linkers are largely unstructured parts of a protein and typically represent intrinsically disordered regions, often mediating protein-protein interactions^56^. This suggests that EVEx do not directly affect the tertiary structure but affect protein functioning and may rewire signalling and regulatory networks in different cell types^57^. Additionally, protein repeats, including WD40 and ankyrin repeat with short repetitive sequence and/or structural motifs were frequently affected. Alternative splicing is known to drive functional and structural diversity in these highly modular domains^58–60^. Protein Kinase-like (PK like) superfamily was enriched highly with adult brain-region specific inclusion of matched cell types (E4), consistent with the known roles of kinases in synaptic plasticity^61^. Additionally, exons with transient developmental exon regulation (E2), which were also lowly recapitulated in human tissues, are enriched for DBL homology domains as well as D-aminoacid aminotransferase-like PLP-dependent enzymes. These superfamilies are associated with cytoskeletal organization and neuronal development and morphogenesis^62^ – processes governing both cell-type identity establishment and differentiation. Finally, adult cell-type specific EVExs (E5), especially those distinguishing neurons from glia, are enriched for Fibronectin type III (Fn3) superfamily. Examples of proteins whose Fn3 domains are affected by EVExs include NRCAM and NFASC, both of which have been associated with neural regulation^63–66.Their^ structure includes immunoglobulin-like (IG-like) domains followed by several Fn3 domain repeats. The presence of Fn3 repeats in tandem with IG domain repeats are not uncommon and found in other proteins associated with regulation of neuronal activities, such as IFN-R-1^67^, IL-6 receptor subunit beta^68^, and Tyrosine-protein kinase^69^ (**Fig S12**). These findings indicate that biological programs defining EVEx inclusion are intrinsically tied to cellular identity and function (**Fig 3j**, Wilcoxon p<2.2e^-16^).

Identified in E2, Exon 3 of the *Jakmip2* gene, annotated as constitutive, is developmentally regulated in hippocampal but not visual cortex excitatory neurons (**Fig 3k**). Similarly, the Testis expressed 9 (*Tex9*) exhibits low brain-region specificity in protein and single-cell RNA data^70, 71^. However, *Tex9* exon 5 is found in E4 and marks brain regions: VIS excitatory neurons have near-constitutive inclusion while other brain regions’ excitatory neurons show as low as 11% (**Fig 3l**). Gene ontology terms associated with neurotransmitter secretion and synaptic potential are enriched in E4, indicating that matched cell types expressing the same gene across multiple brain regions modulate neuronal function using alternative splicing (**Fig 3m**). These findings further underscore the role of alternative exons in conferring developmental, cell-type, and brain-region specificity.

### Adolescent splicing regulation shows a transient increase in brain-region specificity

Given that age alone was not enough to define isoform differences (refer Fig S8), we delineated the dynamic programs of cell-type and brain-region specificity in development. We first considered neuronal cell subtypes from the HIPP and VIS developmental lineages and correlated their Ψ values. Hierarchical clustering of these separated excitatory from inhibitory populations independently of brain region and developmental stage (**Fig 4a**). Surprisingly, pairwise correlations were higher between excitatory types than between inhibitory neuron types (**Fig S13**, two-sided Wilcoxon rank sum test, p=5.099e^-14^). We then considered finer excitatory-cell subtypes (22 cortical types at different ages and eight area-specific hippocampal types at different ages). Similarly, inhibitory cells comprised eight interneuron and four Cajal-Retzius types, while progenitors relating to both lineages comprised NIPCs accounting for eight types, and four granule neuroblast types (GranuleNBs). Clustering of exon inclusion correlation values revealed four main groups, corresponding to: (i) NIPCs, (ii) GranuleNBs, (iii) a group of mature neurons including excitatory and inhibitory subtypes, and (iv) multiple Cajal-Retzius clusters, with a unique cell-type signature (**Fig S14**). Removing Cajal-Retzius subtypes from inhibitory neurons yielded higher intra-inhibitory neuron correlation, but even in their absence, pairwise inhibitory correlations remained lower than pairwise excitatory-cluster correlations. Thus, Cajal-Retzius cells contribute strongly to hippocampal inhibitory neuron diversity, but do not fully account for it (**Fig 4b**). We further investigated timepoints dictating large developmental shifts to examine if these distinguished brain regions. In VIS, excitatory-neuron correlations between adjacent timepoints were lowest at the P21-to-P28 transition with higher values before and after. However, for inhibitory neurons, we found the opposite trend (**Fig 4c**). We correlated exon Ψs between VIS and HIPP for matched cell types. Excitatory neurons showed high HIPP-vs-VIS correlation at P14 and P56, but lower correlations at P21 and especially P28. These observations imply that developmental timelines of excitatory-neuron splicing differ between VIS and HIPP in this period, and brain-region specificity transiently increases. A similar albeit weaker observation was made for inhibitory neurons. The lowest HIPP-vs-VIS correlation occurred at the P21-to-P28 critical developmental period, indicating non-aligned splicing shifts for cortical and hippocampal excitatory and inhibitory neurons (**Fig 4d**). *Bin1,* a synaptic gene implicated in Alzheimer’s disease, exemplifies regional specificity in excitatory neurons and temporal regulation at the P21-P28 transition. Hippocampal excitatory CA neurons express the P1 cerebellar neuronal isoform^70^ at P14 and as the main isoform at P56, which begins to transiently disappear at P21 and is almost entirely absent at P28. At P28, *Bin1* excitatory-neuron isoforms resemble the isoform profile of oligodendrocytes, skipping all six alternative exons. While this is also observed in the visual cortex, the transition is drastic from P21 to P28, marking a brain-region difference between HIPP and VIS (**Fig 4e**). Similar patterns are seen in other disease-relevant genes such as *Mapt* (**Fig S15a-b**). In summary, cell-type and brain-region specificity in isoform expression can be transient for a subset of genes, blurring the lines of cell-type identity as defined by splicing. This raises questions about the effect of the cellular microenvironment on splicing and is an important consideration in functional ramifications especially during development.

**Figure 4.**
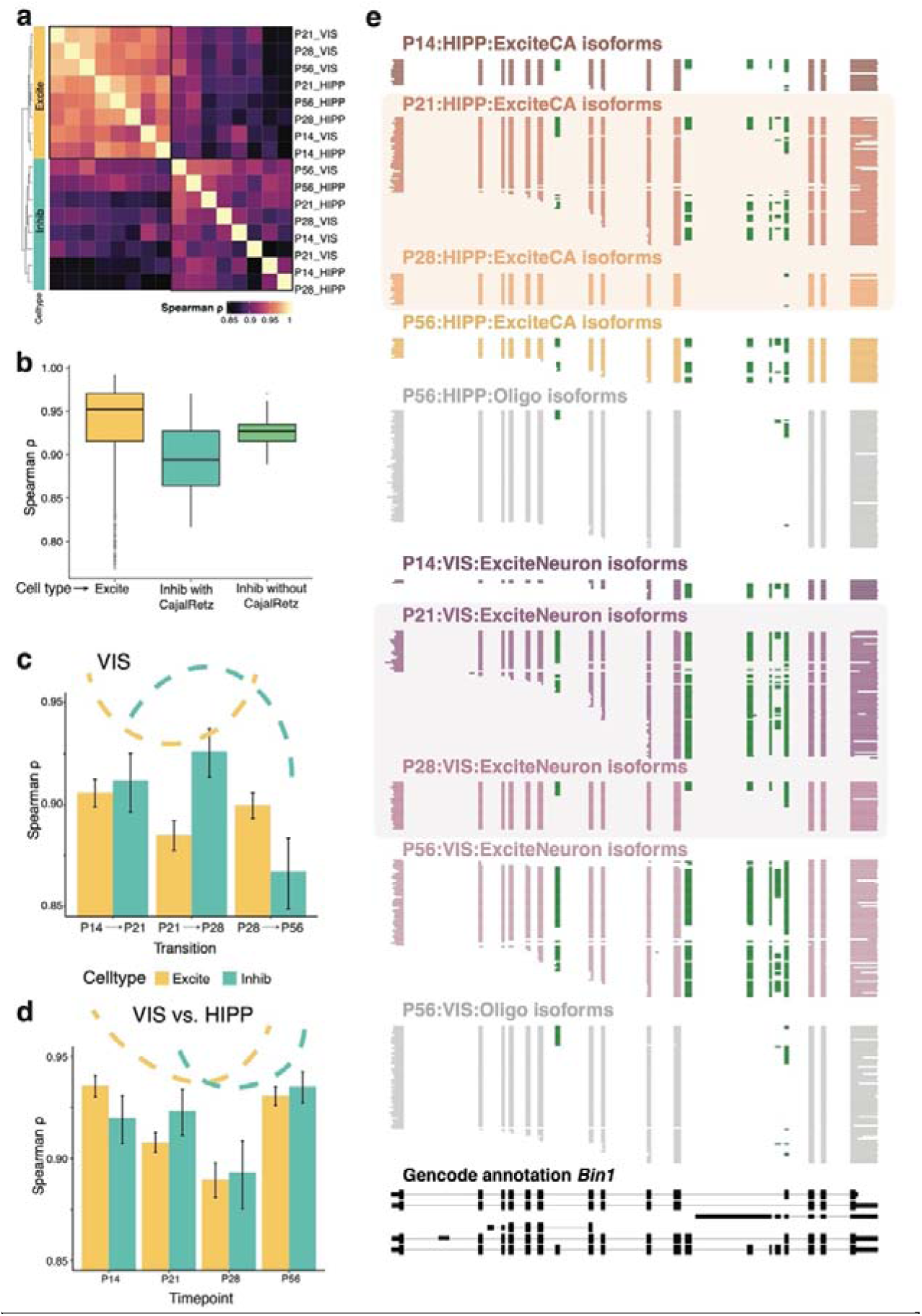
Neuronal exon inclusion changes between mouse visual cortex (VIS) and hippocampus (HIPP) and over time. **(a)** Heatmap of pairwise correlations of exon inclusion (Ψ) for excitatory and inhibitory types. **(b)** Boxplot of pairwise correlations of Ψs for pairs of excitatory subtypes, all inhibitory subtypes, and inhibitory subtypes excluding Cajal-Retzius cells. **(c)** Correlations of exon inclusion for excitatory clusters between neighboring time points in the visual cortex (yellow). Same for inhibitory clusters (green) **(d)** Correlations of exon inclusion for excitatory clusters at each time point between visual cortex and hippocampus (yellow). Same for inhibitory clusters (green) **(e)** Depiction of cell type-resolved single-cell long reads for the *Bin1* gene in HIPP and VIS excitatory neurons. Each line represents one individual cDNA molecule, and blocks are colored by cell type and timepoint. Green represents alternative exons. Grey blocks indicate oligodendrocyte populations at P56. Bottom black track: Gencode annotation

### A splicing switch in oligodendrocyte maturation occurs after a gene-expression defined split from astrocytes

Similar to the previous analysis of neuronal types, using timepoint-resolved visual cortex and hippocampal data for astrocyte and oligodendrocyte subtypes, we determined exon inclusion levels for 35 corresponding clusters. We represented clustering based on pairwise correlations of these cluster-level exon inclusion profiles in a heatmap. Surprisingly, the first split in the dendrogram separated astrocytes and all OPC clusters -regardless of age-from all committed and mature oligodendrocytes (**Fig 5a**). This is in stark contrast to similar analysis based on gene expression profiles for each cluster, which groups all oligodendrocyte-lineage clusters together – but well separated from astrocytes (**Fig 5b**). Consistent with these observations, a pseudotime trajectory analysis^72^ using single-cell isoform expression data (Methods) with a starting point defined at OPCs revealed two trajectories, one towards astrocytes and one along the oligodendrocyte lineage. Of note, the trajectory from OPCs to astrocytes likely does not represent a maturation pattern, but rather the fact that mathematically OPCs are close to astrocytes in terms of splicing patterns (**Fig 5c**). Taken together, these analyses support divergent cell-group similarities observable in splicing and 3’ gene expression patterns which yet again motivate cell-type identity in part defined by splicing (**Fig 5d**).

**Figure 5.**
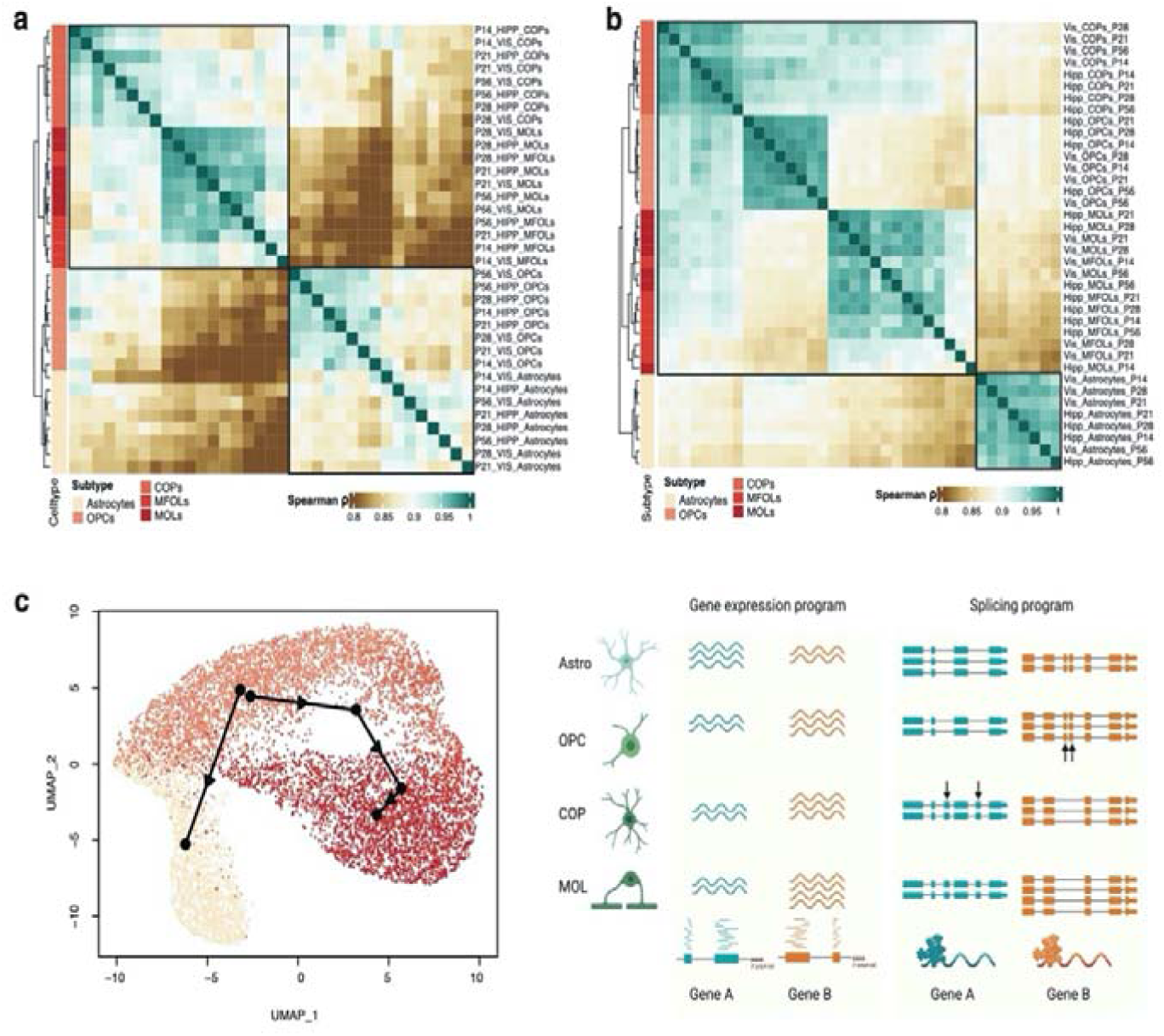
Exon inclusion patterns in glial subtypes suggest an ordered molecular cascade. **(a)**. Heatmap based on pairwise correlations of exon inclusion patterns for astrocyte and oligodendrocyte-lineage cells. **(b)** Similar heatmap based on pairwise gene expression values. **(c)** Slingshot trajectory of glial cells using exon inclusion values. **(d)** Model depiction summarizing findings of previous panels: Subtypes in the oligodendrocyte lineage have similar gene expression patterns. However, a switch in splicing patterns occurs after OPCs have matured to COPs. Arrows represent alternative exons.

### Fluctuating patterns of exon variability between cell types in a critical developmental period

We hypothesized that EVExs with temporal regulation (E1-E2, Fig 3e) were related to the correlation drop around P28 (Fig 4c-d). We focused on 1,072 exons that substantially change exon variability between timepoints (**Fig S16**, Methods). Three transitions, each of (i) increasing, (ii) decreasing, or (iii) constant variability define 3^3^=27 potential patterns – but after removing invariable exons (G0), only 9 were frequent (denoted as G1-G9, **Fig 6a**). For genes containing multiple alternative exons, 36% had all exons exhibiting fluctuations in a single pattern (n = 106 HIPP, n = 115 VIS) while 64% had exons in two or more patterns (n = 195 HIPP, n = 204 VIS). Interestingly, we found that pairs of neighboring exons were frequently included or skipped together (**Fig S17a**). As hypothesized, the P21-P28 transition consistently exhibited drastic shifts in exon variability (**Fig 6b**, **Fig S18-S19,** Bernoulli p-value 1.89e-06). The exon-inclusion variability between cell types had the highest standard deviation at P28 in both HIPP and VIS, and we found that pairs of exons were tightly coordinated at this timepoint (**Fig S17b**). These observations further suggest that the difference between cell types can either disappear or be enhanced during this critical time period (**Fig S20**).

**Figure 6:**
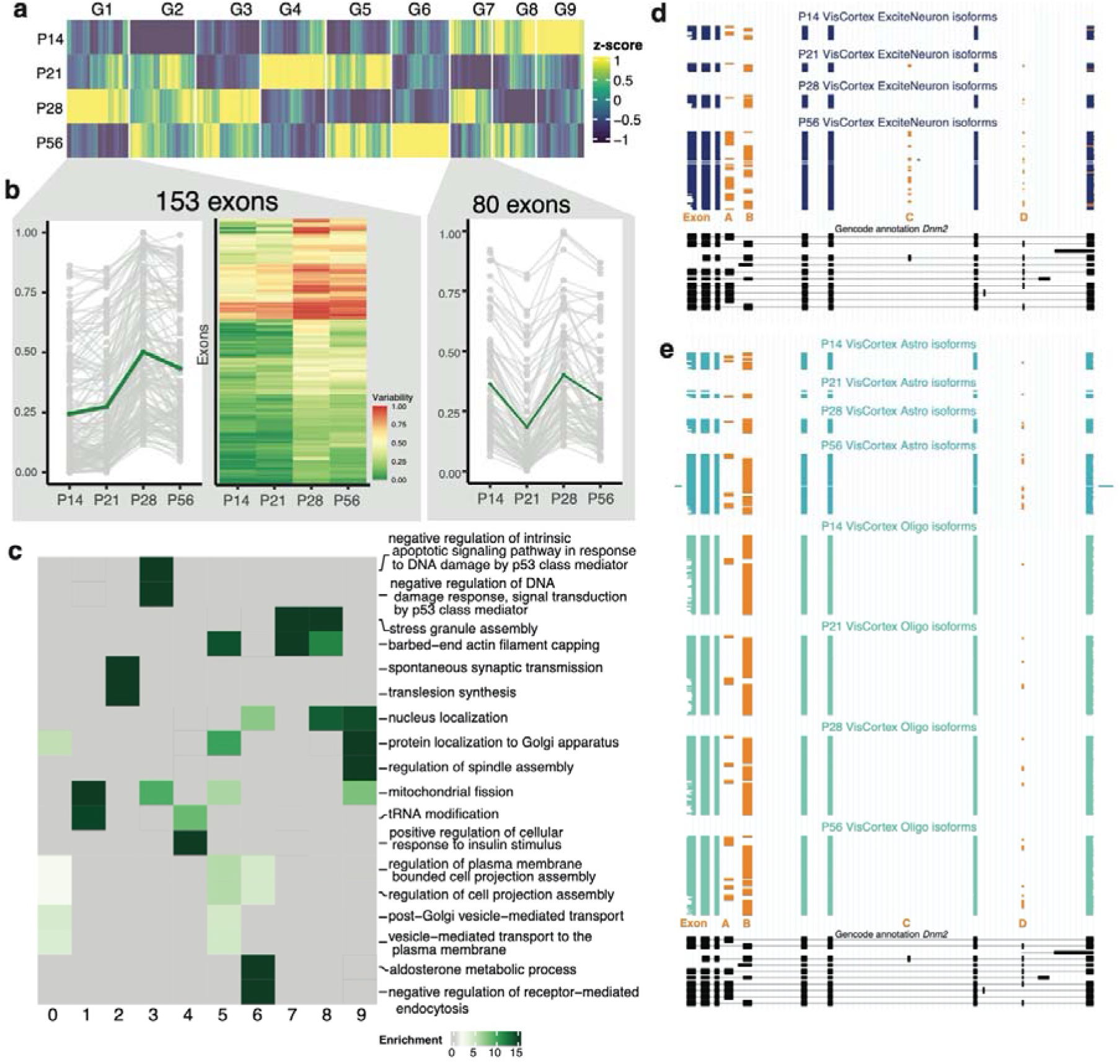
Developmental exon regulation reveals convergent and divergent patterns of exon variability. **(a)** Heatmap of z-normalized exon variability between the four major cell types across development **(b)** Line plot of values of exon variability for individual genes in groups 3 (left) and 7 (right) for VIS. Heatmap of exon variability for group 7 in VIS. Some show lower changes while others exhibit drastic differences **(c)** Heatmap of gene ontology (GO) enrichment values for highly enriched sets of genes contributing to the 9 groups from (a). **(d)** Depiction of timepoint-resolved single-cell long reads from visual cortex excitatory neurons for the *Dnm2* gene. Each line represents one individual cDNA molecule. Alternative exons denoted in orange and marked A through D. **(e)** Same as in (d) but for astrocyte (teal) and oligodendrocyte (sea-green) clusters. Bottom black track: Gencode annotation

Genes unique to each of the G1-G9 patterns were largely non-overlapping in function (**Fig S21**). Of note, synaptic vesicle budding and transport was associated with genes exhibiting an exon variability increase at P28 (G1) whereas actin filament capping and functions associated with cell projection were associated with the aforementioned G5 where we see an increase in exon variability at P21 and P56. Both observations likely indicate a developmentally regulated neuronal versus glial split (**Fig S21, Fig 6c**). The Dynamin2 (*Dnm2*) gene is ubiquitously expressed, has known roles in intracellular membrane trafficking and cytoskeleton organization, and has previously been associated with neurological diseases^73^. In VIS, *Dnm2* has exons in G5 and G8 and therefore exemplifies repetitive developmental changes in cell-type variability. The ScisorWiz^74^ plot shows *Dnm2* leveraging four alternative exons to produce 6 complex isoforms. In excitatory neurons at P14, two mutually exclusive exons (Exons A,B) define two major isoforms, the second of which dominates at P21 and P28. At P56 however, the first exon gets upregulated to again resemble the pattern at P14. However, two additional exons render the isoform landscape more complex, where Exon C is only visible in P56 excitatory neurons while Exon D starts upregulating its expression at P28 (**Fig 6d**). Exons A and B in glia at all timepoints resemble the splicing patterns of excitatory neurons at P21 and P28, but not at P14 and P56. In contrast to excitatory neurons, exon C is not seen in glia in the ages and region examined, while exon D is expressed uniformly starting at P28 (**Fig 6e**). *Dnm2* not only varies across the cell-type and developmental axes but also shows differing patterns of isoform regulation between VIS and HIPP (compare Fig 6d-e to **Fig S22**). This example highlights how cell types leverage inclusion patterns of multiple exons in tandem for developmental and brain-region specific specialization.

## Discussion

Isoform expression underlies physiological differences in brain regions and cell types and is altered in development and evolution. Developmental regulation of alternative splicing further encapsulates cellular differentiation and maturation. However, while a large body of research had recognized that isoform changes occur during brain development and evolution, a unified view of these dimensions of anatomic regional and cell type differences over time remained elusive.

We found that cell types vary extensively in their splicing patterns across multiple investigated axes. While the number of exons modulating their variability between cell types within a single brain region exceeds that of the same cell type between brain regions, we find compelling evidence for the latter, involving neurotransmitter secretion and regulation as well synaptic regulation. Thus, while the establishment of cell-type identity is essential for brain function and regional specialization, cell types that are considered homogenous and are ubiquitously present in brain tissue display specialized splicing patterns depending on their region of origin. This is particularly true for astrocytes in the thalamus and the cerebellum, which differ not only in their exon inclusion patterns, but also in their TSS and PolyA-site usage, rendering the mRNA isoform landscape more complex than previously imagined.

We parsed through this complex landscape and defined groups of exons with extreme variability (EVExs) seen across brain regions, development, cell types, as well as in combinations thereof. These exon classes exhibit key differences in intrinsic properties such as length, protein coding capacity, and protein domain architecture. Therefore, distinct programs of exon variability correlate with functional consequences of the protein product. This still leaves some questions unanswered such as the precise definition of regulatory elements governing the establishment of these variability classes and their conservation in human. Nonetheless, we found that adult cell-type variability is largely recapitulated in human single-nuclei long-read data. Therefore, many of the mouse results can be extrapolated to large cell-type specific splicing changes in perturbed and disease conditions to human endeavors.

Importantly, we find that the same cell type traced along the developmental axis exhibits more fluctuation in its splicing patterns than it does across brain regions in adulthood. Synapse formation, axon guidance, and general neural network formation induce a temporal heterogeneity in splicing patterns within cell populations that are attenuated in adulthood. This observation is further strengthened by nine patterns of developmental variability that we identify, a majority of which involve fluctuations during the critical developmental period^75^ of murine adolescence (P21-P28) in the visual cortex and hippocampus. We repeatedly find that exons that are cell-type specific at a developmental timepoint can transiently change inclusion status and lose their cell-type specificity. Fundamental genes, such as *Bin1* and *Mapt* show such transient cell-type specificity in isoform expression which suggests a highly sophisticated developmental splicing program. Additionally, our data reveal the precise timing of splicing switches in maturation programs, especially on the oligodendrocyte lineage after the split from astrocytes. This adds a layer of subtlety to studies of isoform expression over development and justifies the need for simultaneous recording of gene expression and splicing to define the relationships between cell types and across brain regions.

Taken together, we present a comprehensive single-cell investigation of alternative isoform usage in mouse and human brain and the dynamics of isoform expression variation across anatomical structures and development.

## Acknowledgments

This work is supported through the BRAIN Initiative Cell Census Network grant 1RF1MH121267-01 to H.U.T. We thank Jason McCormick and Tomas Baumgartner from Weill Cornell Medicine Flow Cytometry Core Facility for FACS assistance, and Dong Xu, Xing Wang, Adrian Tan, and Jenny Xiang from the Genomics Resources Core Facility for performing RNA sequencing. We thank Dr. Christopher Mason for use of his PromethION machine. We also thank Weill Cornell Medicine Scientific Computing Unit (SCU) for use of their computational resources. H.U.T. is additionally supported by NIGMS grant 1R01GM135247-01, NIDA grant U01 DA053625-01, and by the Feil Family Foundation. M.E.R is supported by the National Institute of Health (NIH) grants 1R01NS105477, P01HD067244, U54NS11717, and by the Feil Family Foundation. D.K. is supported by National Library of Medicine of the National Institutes of grant R01LM014017. T.A.M. is supported by NIH grants DA08259 and HL136520. L.C.N. is supported in part by the NIMH, NIDA, NINDS, NIDDK, NHLBI, and NIAID under award number: UM1AI164599 and by NIDA under award number U01 DA53625 (to L.C.N., H.T., And T.M). M.D is supported by NIH/NHGRI U41HG007234. E.D.J and O.F. are supported by funds from HHMI.

## Author Contributions

A.J. and H.U.T. conceived the project and designed experiments and computational analyses. W.H., B.Z., J.B., performed experiments. A.J., O.N., and M.D. conducted computational analyses. E.J., G.S., D.K., M.E.R., H.U.T. supervised the project. T.A.M, L.C.N, O.F., contributed key reagents. A.J., H.U.T. wrote the manuscript. All authors participated in the review and editing of the manuscript.

## Methods

### Ethics statement

All experiments were conducted in accordance with relevant NIH guidelines and regulations, related to the Care and Use of Laboratory Animals tissue. Animal procedures were performed according to protocols approved by the Research Animal Resource Center at Weill Cornell Medicine.

### ScISOr-Seq2: Single cell isoform RNA sequencing from adolescent and adult mouse brain

#### Tissue acquisition (mouse)

C57BL/6NTac mice were perfused with 25mL ice-cold and carbogen treated 1X partial-sucrose cutting solution containing 5 µg/mL actinomycin D. The remaining 1X partial sucrose cutting solution and EBSS was oxygenated. Dissection of specific brain regions was conducted by using the Mouse 3D Coronal sections from Allen Brain Atlas as a reference map for coordinates. The brain slices were collected on Vibratome (Leica) at the thickness of 300 µm/slice and kept in ice-cold 1 X partial sucrose cutting solution. For the hippocampus, 8∼10 mouse coronal slices (300µm) were collected from the caudal region of the brain after removing of the cerebellum. The hippocampus region was dissected out based on the Mouse Coronal Sections (images 62∼89). Note: each image section on Allen Brain Atlas is spaced at 100 µm interval. 8∼10 slices can cover almost the whole hippocampus region. Visual cortex was collected based on the images 79∼100. The first 1 or 2 slices from the caudal region of the brain were discarded and subsequently 5∼6 continuous slices were collected. For striatum, 5∼6 mouse coronal slices were collected from the rostral region of the brain after removing the olfactory bulb. The striatum was dissected out based on the Mouse coronal Sections (images 39∼59). Dissecting of cerebellum does not require any vibratome sectioning. The cerebellum was dissected based on the location and structure with forceps and minced into small pieces with scalpel. Slices were transferred to slicing chamber with bubbling 1X partial sucrose-containing small molecule mix B at RT – and slices were allowed to recover for ∼30 minutes. 1X cutting solution: 93 mM N-Methyl-D-glucamine (Acros Organics, AC126841000), 2.5 mM KCl (Sigma, 44675), 1.2 mM NaH2PO4 (Sigma, S5011), 30 mM NaHCO3 (Sigma, S5761), 20 mM Hepes (Gibco, 15630106), 25 mM glucose (Sigma, G7021), 5 mM sodium ascorbate (Sigma, A4034), 2 mM thiourea (Alfa Aesar, AAA1282822), 3 mM sodium pyruvate (Gibco, 11360070), 10 mM N-Acetyl-L-Cysteine (Alfa Aesar, AAA1540914), 0.5 mM CaCl2 (Sigma, 223506), 10 mM MgSO4 (Sigma, M2643), pH7.2-7.4.

#### Single cell disassociation (mouse)

Tissue sections were dissociated with modification from a previous protocol from the Smit lab at VU, Netherlands: Regions of interest were dissected on a Sylgard coated plate with dark background, in 1-2 mL carbogen treated cutting solution. Tissue pieces were transferred to 5 mL of 2 mg/mL activated papain (Wortington, LK003150) then incubated for 15-25 min at 37 °C with gentle mixing. After the incubation, tissue was cut into tiny pieces and then gently triturated 15–20 times using Large to small size asteur pipettes until no obvious chunks can be observed. Pasteur pipettes with different openings sizes (Large: 0.6–0.7 cm, middle: 0.3–0.4 cm, small: 0.15-0.2 cm) were created by flame polishing disposable glass pasteur pipettes (ThermoFisher) and assembled with rubber bulbs. After undissociated tissue chunks settled down, supernatant was taken and then filtered using a 30 μm cell strainer (Miltenyi Biotec, 130-041-407) into a nuclease free collection tube. The supernatant was then centrifuged at 300-400 rcf for 5 min at RT. After discarding the supernatant, the cell pellet was resuspended in 3 ml 10% ovomucoid protease inhibitor solution (150 µL DNase I, 300 µL ovomucoid protease inhibitor solution and 2.55 mL EBSS, Wortington, LK003150). Next, the cell suspension was slowly and gently added to the top layer of 5 mL ovomucoid protease inhibitor solution (Wortington, LK003150) without interfering with the bottom layer. The cells were spun down after centrifugation at 70-100 g for 6 min at RT. After removing all the supernatant, cells were suspended in 1 mL FACS buffer: 1X HBSS (Gibco, 14175079) containing 0.2% BSA (Thermo Scientific, 37525), 25mM glucose (Sigma, G7021), 3mM sodium pyruvate (Gibco, 11360070), and 0.2U/µL RNase inhibitor (Ambion,AM2682). After incubation for 15 min in FACS buffer with 0.1 µg/mL DAPI (Sigma, D9542), viable cells were collected as DAPI negative population using Sony MA900 sorter with FlowJo version 10 software. Sorted viable cells were centrifuged down and subsequently diluted to 1000-1500 cells/μL in FACS buffer for capture on the 10x Genomics Chromium controller.

### SnISOr-Seq: Single nucleus isoform sequencing in frozen human tissue

#### Sample acquisition (human)

Hippocampal samples for SnISOr-Seq: Six healthy human brain samples (hippocampus, 3M, 3F, Table S5) used for this study were requested through the NIH NeuroBioBank and obtained from the University of Maryland Brain and Tissue Bank according to Institutional Review Board-approved protocols. No subjects had pre-existing neurodegenerative or neurological disease. Tissues were flash frozen and kept at −80°C until processing.

#### Single-nuclei isolation (human)

Single nuclei suspension was isolated from fresh-frozen human brain samples using the SnISOr-Seq^1^ protocol. In brief: Approximately 30 mg of frozen tissue of each sample was dissected in a sterile dish on dry ice and transferred to a 2 mL glass tube containing 1.5 mL nuclei pure lysis buffer (MilliporeSigma, catalog no. L9286) on ice. Tissue was completely minced and homogenized to nuclei suspension by sample grinding with Dounce homogenizers (Millipore Sigma, catalog no. D8938-1SET) with 20 strokes with pestle A and 18 strokes with pestle B. The nuclei suspension was filtered by loading through a 35 µm diameter filter and followed by centrifuging 5 min at 600 g and 4℃. The nuclei pellet was collected and washed with cold wash buffer, which consisted of the following reagents: 1X PBS (Corning, catalog no. 46-013-CM), 20 mM DTT (Thermo Fisher Scientific, catalog no. P2325), 1%BSA (NEB, catalog no. B9000S), 0.2U/µL RNase inhibitor (Ambion, catalog no. AM2682) for three times. After removing the supernatant from the last wash, the nuclei were resuspended in 1 mL of 0.5 µg/mL DAPI (Millipore Sigma, catalog no. D9542) containing wash buffer to stain for 15 min. The nuclei suspension was prepared for sorting by filtering cell aggregates and particles out with a diameter of 35 µm. After removing myelin and fractured nuclei by sorting, the nuclei were collected by centrifuging 5 min at 600 g and 4℃, then resuspended in wash buffer to reach a final concentration of 1×10e6 nuclei/mL after counting in trypan blue (Thermo Fisher Scientific, catalog no. T10282) using a Countess II cell counter (Thermo Fisher Scientific, catalog no. A27977).

#### 10x 3’ library preparation and sequencing

A single cell/nuclei suspension containing 10,000 cells/nuclei was loaded on a Chromium Single Cell B Chip (10x Genomics, catalog no. 1000154) as follows: 75 µL of master mix + nuclei suspension was loaded into the row labeled 1, 40 µL of Chromium Single Cell 3ʹ Gel Beads (10x Genomics, catalog no. PN-1000093) into the row labeled 2 and 280 µL of Partitioning Oil (10x Genomics, catalog no. 2000190) into the row labeled 3. This was followed by GEM generation and barcoding, post GEM-RT cleanup and cDNA amplification. Then 100 ng purified cDNA derived from 12 cycles of cDNA amplification was used for 3ʹ Gene Expression Library Construction by using Chromium Single Cell 3ʹ GEM, Library & Gel Bead Kit v3 (10x Genomics, catalog no. 1000092) according to the manufacturer’s manual (10x Genomics, catalog no. CG000183 Rev C). The barcoded short-read libraries were measured using a Qubit 2.0 with a Qubit dsDNA HS assay kit (Invitrogen, catalog no. Q32854) and the quality of the libraries was assessed on a Fragment analyzer (Agilent) using a high-sensitivity NGS Fragment Kit (1-6000bp) (Agilent, catalog no. DNF-474-0500). Sequencing libraries were loaded on an Illumina NovaSeq6000 with PE 2 x 50 paired-end kits by using the following read length: 28 cycles Read1, 8 cycles i7 index and 91 cycles Read2.

### Linear/asymmetric PCR and exome capture (LAP-CAP)

#### Linear/asymmetric PCR to remove non-barcoded cDNA

The first round PCR protocol (95 °C for 3 min, 12 cycles of 98 °C for 20 s, 64°C for 30 s and 72 °C for 60 s) was performed by applying 12 cycles of linear/asymmetric amplification to preferentially amplify one strand of the cDNA template (30 ng of cDNA generated by using 10x Genomics Chromium Single Cell 3ʹ GEM kit) with primer ‘Partial Read1’, and then the product was purified with 0.8× SPRIselect beads (Beckman Coulter, B23318) and washed twice with 80% ethanol. The second round PCR is performed by applying six cycles of exponential amplification under the same condition with forward primer ‘Partial Read1’ and reverse primer ‘Partial TSO’, and then the product was purified with 0.6× SPRIselect beads and washed twice with 80% ethanol and eluted in 30 µl of buffer EB (Qiagen, 19086). Sequences of primers: Partial Read1 (5′-CTACACGACGCTCTTCCGATCT-3′) and Partial TSO (5′-AAGCAGTGGTATCAACGCAGAGTACAT-3′). KAPA HiFi HotStart PCR Ready Mix (2×) (Roche, KK2601) was used as polymerase for all the PCR amplification steps in this paper, except for the 10x Genomics 3′ library construction part.

#### Exome capture to enrich for spliced cDNA

Exome enrichment was applied to the cDNA purified from the previous step by using probe kit SSELXT Human All Exon V8 (Agilent, 5191-6879) for human samples, or SureSelectXT Mouse All Exon (Agilent, 5190-4641) for mouse samples. The reagent kit SureSelectXT HSQ (Agilent, G9611A) was used according to the manufacturer’s manual. First, the block oligo mix was made by mixing an equal amount (1 µl of each per reaction) of primers Partial Read1 (5′-CTACACGACGCTCTTCCGATCT-3′) and Partial TSO (5′-AAGCAGTGGTATCAACGCAGAGTACAT-3′) with the concentration of 200 ng/µL (IDT), resulting in 100 ng/µL. Next, 5 µl of 100 ng/µLcDNA diluted from the previous step was combined with 2 µL of block mix and 2 µL of nuclease free water (NEB, AM9937), and then the cDNA block oligo mix was incubated on a thermocycler under the following conditions to allow block oligo mix to bind to the 5′ end and the 3′ end of the cDNA molecule: 95 °C for 5 min, 65 °C for 5 min and 65 °C on hold. For the next step, the hybridization mix was prepared by combining 20 mL of SureSelect Hyb1, 0.8 ml of SureSelect Hyb2, 8.0 mL of SureSelect Hyb3 and 10.4 mL of SureSelect Hyb4 and kept at room temperature. Once the reaction reached to 65 °C on hold, 5 µL of probe, 1.5 µL of nuclease-free water, 0.5 µL of 1:4 diluted RNase Block and 13 µL of the hybridization mix were added to the cDNA block oligo mix and incubated for 24 h at 65 °C. When the incubation reached the end, the hybridization reaction was transferred to room temperature. Simultaneously, an aliquot of 75 µL of M-270 Streptavidin Dynabeads (Thermo Fisher Scientific, 65305) were prepared by washing three times and resuspended with 200 µl of binding buffer. Next, the hybridization reaction was mixed with all the M-270 Dynabeads and placed on a Hula mixer for 30 min at room temperature. During the incubation, 600 µL of wash buffer 2 (WB2) was transferred to three wells of a 0.2 mL PCR tube and incubated in a thermocycler on hold at 65 °C. After the 30-min incubation, the buffer was replaced with 200 µL of wash buffer 1 (WB1). Then, the tube containing the hybridization product bound to M-270 Dynabeads was put back into the Hula mixer for another 15-min incubation with low speed. Next, the WB1 was replaced with WB2, and the tube was transferred to the thermocycler for the next round of incubation. Overall, the hybridization product bound to M-270 Dynabeads was incubated in WB2 for 30 min at 65 °C, and the buffer was replaced with fresh pre-heated WB2 every 10 min. When the incubation was over, WB2 was removed, and the beads were resuspended in 18 µL of nuclease-free water and stored at 4 °C. Next, the spliced cDNA, which bound with the M-270 Dynabeads, was amplified with primers Partial Read1 and Partial TSO by using the following PCR protocol: 95 °C for 3 min, 12 cycles of 98 °C for 20 s, 64 °C for 60 s and 72 °C for 3 min. The amplified spliced cDNA was isolated from M-270 beads as supernatant and followed by a purification with 0.8× SPRIselect beads.

**Library preparation for PacBio**

HiFi SMRTbell libraries were constructed according to the manufacturer’s manual by using SMRTbell Express Template Prep Kit 2.0 (PacBio, 100-938-900). For all samples, ∼500 ng of cDNA obtained by performing LAP-CAP from the previous step was used for library preparation. The library construction includes DNA damage repair (37 °C for 30 min), end-repair/A-tailing (20 °C for 30 min and 65 °C for 30 min), adaptor ligation (20 °C for 60 min) and purification with 0.6× SPRIselect beads.

#### Library preparation for ONT

For all samples, ∼75 fmol cDNA was processed with LAP-CAP underwent ONT library construction by using the Ligation Sequencing Kit (ONT, SQK-LSK110), according to the manufacturer’s protocol (Nanopore Protocol, Amplicons by Ligation, version ACDE_9110_v110_revC_10Nov2020). The ONT library was loaded onto a PromethION sequencer by using PromethION Flow Cell (ONT, FLO-PRO002) and sequenced for 72 h. Base-calling was performed with Guppy by setting the base quality score >7.

#### Short read assignment of cell types (mouse)

Fastq files were obtained from the Illumina sequencing reads by running bcl2fastq. Gene x cell matrices processed with cellranger V3.1.0 were loaded into Seurat V3.2.3^2^ and preprocessed individually using cutoffs described in Table S4. After filtering for high quality cells, they were scaled and normalized using default parameters and clustered using the Louvain algorithm. Doublet clusters were discarded. Subsequently, all samples from the hippocampal developmental lineage were processed together, as were the samples from the visual cortex lineage. After combining the data without any integration approaches and using the Seurat merge function, the data was scaled, normalized, and variable genes identified. Integration of data to control for sample-specific batch effects was done using Harmony. Cell types were assigned using marker genes in three levels of granularity – broad, cell type, and cell subtype. These were then assigned to each single cell along with the information on replicate, brain region, age. For the spatial axis i.e., the cerebellum, striatum, and thalamus, the two replicates were integrated with Harmony^3^ before assigning cell types in the same three levels of granularity as above. Finally, the entire dataset was merged together into a single object for visualization purposes and to obtain summary statistics in Fig 1. This was also done using harmony while controlling for region-specific differences in gene expression.

#### Short read assignment of cell types from human hippocampal data

Fastq files were obtained from the Illumina sequencing reads by running bcl2fastq. Gene x cell matrices processed with cellranger v3.1.0 were loaded into Seurat 3.2.2 and preprocessed individually. After filtering for high quality cells, they were scaled and normalized using default parameters and clustered using the Louvain algorithm. Doublet clusters were discarded. Subsequently, all samples were processed together. After combining the data without any integration approaches and using the Seurat merge function, the data was scaled, normalized, and variable genes identified. Integration of data to control for sample-specific batch effects was done using Harmony ^3^. Cell types were assigned using marker genes in three levels of granularity – broad, cell type, and cell subtype. These were then assigned to each single cell along with the information on sample id.

#### Generation of PacBio circular consensus reads

Using the default SMRT-Link (v8.0.0.78867) parameters, we performed circular consensus sequencing (CCS, 8.0.0.80529) with IsoSeq3 with the following modified parameters: maximum subread length 14,000 bp, minimum subread length 10 bp, and minimum number of passes 3.

#### Preprocessing of long read sequencing data with PacBio

Subread fastq files were obtained, CCS performed using parameters described above. Data was processed using the scisorseqr^4^ pipeline by first aligning to the genome using STARlong (v2.7.0). Cellular barcodes were assigned using the cell type and sample information from the short read analysis as input to the GetBarcodes() function. Subsequently, uniquely mapped, spliced, barcoded reads were obtained using the MapAndFilter() and InfoPerLongRead() functions in scisorseqr.

#### Preprocessing of long read sequencing data with ONT

Reads were basecalled using MinKNOW Core (v 4.0.5), Bream (v6.0.10), and guppy (v4.0.11) on the PromethION machine. Reads were aligned using minimap2^5^ (v2.17-r943-dirty) and data was preprocessed using the scisorseqr (v0.1.9)^4^ package. Cell barcodes were assigned using the cell type and sample information from the short read analysis.

#### PacBio transcript assignment using IsoQuant

IsoQuant (v2.3.0) was run using default PacBio parameters on an aggregate of all barcoded PacBio reads with GENCODE v21 as annotation. Multi-exonic transcripts classified as “novel in catalog” were then used to create an enhanced annotation

#### ONT transcript assignment using IsoQuant

This enhanced annotation gtf file was used on each of the ONT samples to correct incorrectly assigned splice sites in multi-exonic barcoded reads. Isoquant (v3.1) was run using default parameters for ONT data. These corrected splice sites were then re-assigned to reads in the AllInfo file, which was then filtered for unique UMIs, and used as input in subsequent analysis.

#### Obtaining exon counts using corrected splice sites

Using all exons appearing as internal exon in a read, we calculated:

1. The number of long-read molecules containing this exon with identity of both splice sites: *X_in_*
2. The number of long-read molecules assigned to the same gene as the exon, which skipped the exon and >=50 bases on both sides: *X_out_*
3. The number of long-read molecules supporting the acceptor of the exon and ending on the exon: *X_acc_In_*
4. The number of long-read molecules supporting the donor of the exon and ending on the exon: *X_don_In_*
5. The number of long-read molecules overlapping the exon: *X_tot_*

Non-annotated exons with one or two annotated splice sites, ≥70 bases of non-exonic (in the annotation) bases, were excluded as intron-retention events or alternative acceptors/donors.

We then calculated

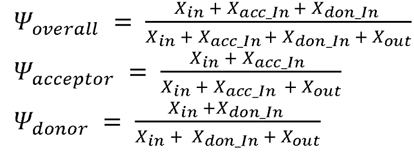

If

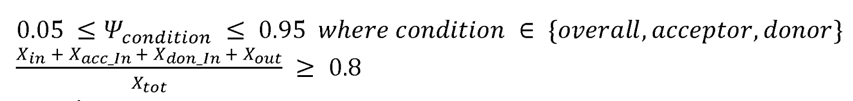

the exon was kept.

We then calculated the *Ѱ_overall_* for each cell type from all long-read UMIs for that cell type if and only if. *X_tot_* 10 for the exon and cell type in question. Otherwise, *Ѱ_overall_* for the exon and cell type was set to “NA”.

#### Obtaining full length isoform variability across three axes

To find regional variability, we first calculate the percent inclusion (PI, Π) for each isoform in a region by summing the inclusion values over all cell subtypes and time points for that region. If at least two regions have Π values, we obtain an *isoform x region* matrix of Π values. Then we define the variability per isoform as the max(Π) – min(Π). The brain region contributing the most to the region-specific variability is recorded as having the Π with the highest divergence from the median Π value.

The same procedure was carried out to calculate age and subtype variability of a given cell type. More formally,

For a cell type (CT), all possible clusters can be represented as a combination of brain region (a1), age (a2), and subtype (a3).

Therefore for

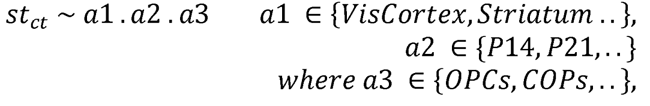

Per gene and cell type (CT), a matrix (G) can be constructed with *i* rows containing the isoforms of the gene and *j* columns containing the cell clusters. So,

*G ~ g_i,j_ where j e st_ct_ and i isofom*

So for a brain region (x), a subset of the matrix can be obtained containing only the subtypes originating from that brain region

*G ~ g_i,j_ where j’ a1 = x. a2.a3*

Thus, for a cell type (CT), the brain region variability for isoform (i) is defined as

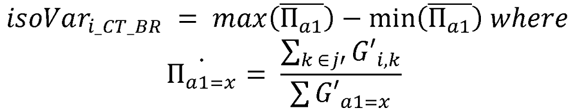

Similarly, the age variability is defined as

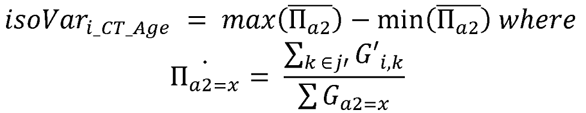

And the subtype variability is defined as

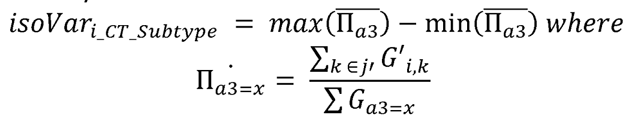

For each isoform, we thus had a raw value for age, region, and subtype variability. If any of these values were at least 0.1 i.e., exhibiting at least a 10% change in isoform usage across an investigated axis, then the values were normalized to add up to 1 and represented in the ternary plot. For values where the normalized regional variability was greater than 0.5 but the age and subtype variability was less than 0.5, then the isoform was considered to be region-specific for a particular cell type.

#### Obtaining full length isoform variability across three axes in pseudo-bulk

A similar analysis to the one above was performed, except that the cell subtype axis was replaced by a cell type axis. Therefore, to find regional variability, we first calculate Π for each isoform in a region by summing the inclusion values over all cell types and time points for that region, and the variability values is obtained by subtracting the min(Π) from the max(Π). Similarly, age variability is calculated by summing the inclusion values of all cell types and regions for a given timepoint. Cell type variability is calculated by summing the inclusion values of all regions and timepoints for a given cell type and the variability is reported as the max(Π) – min(Π) across all cell types.

#### Highly variable gene identification

We filtered the matrix of variability along the three axes for isoforms that had a raw variability value of at least 0.25 along one of the axes. Furthermore, if the isoforms were present in the center trangle, we filtered the isoforms that had a variability of 0.25 in all three axes. These were then clustered together using hierarchical clustering, and plotted as a heatmap per cell type with Complex Heatmap^6^.

#### Differential isoform expression for one brain region vs all

We obtained transcript counts per cell type from IsoQuant as described above. For each major cell type, we obtained an *isoform X region* matrix of counts. To perform differential isoform expression (DIE) analysis, we used the two-sample framework using χ^2^ tests of abundance as described in scisorseqr ^4^. In brief: For each brain region, we aggregated counts from the other four brain regions as a background and performed DIE tests. For each differential abundance test between two categories, genes were filtered out as ‘untestable’ if reads did not reach sufficient depth (25 reads/gene category). For genes with sufficient depth, a maximum of an 11 × 2 matrix of counts denoting *isoform x category* was constructed with the first ten rows corresponding up to the first ten isoforms, and the last row comprised of collapsed counts from all the other isoforms (if any). P-values from a χ^2^ test were reported per gene, along with a ΔΠ value per gene. The ΔΠ was constructed as the sum of change in percent isoform (Π) of the top two isoforms in either positive or negative direction. After these numbers were reported for all testable genes for a comparison, the Benjamini Hochberg (BH) correction for multiple testing with a false discovery rate of 5% was applied to return a corrected p-value. If this FDR p-value was ≤0.05 and the Π was more than 0.1 in one direction, i.e., the change in percent inclusion of one or two isoforms was more than 10% from one category to the other, then the gene was considered to be significantly differentially spliced.

#### Differential TSS, PolyA expression for one brain region vs all

For each sequenced long read, we assigned a known CAGE peak if the start of the read fell within 50 bp of an annotated peak. Similarly, we assigned a PolyA site to a read if it fell within 50 bp of a known site. Counts for each known TSS and PolyA site were obtained per cell type, and reads where a TSS / PolyA site could not be assigned were discarded. Counts for brain region comparison were aggregated in a similar fashion to the full-length transcript analysis described above to obtain an *n x 2* matrix. DIE was performed using scisorseqr, and the ΔΠ was recorded for each gene.

#### Identifying highly variable exons (hVEx)

We first obtained a matrix of Ψ values for all major cell types i.e., astrocytes, oligodendrocytes, excitatory neurons, and inhibitory neurons for each of the 11 samples by summing the counts over all replicates. This yielded 44 triads defined by the age, region, and cell type of origin. Then we considered four lineages to calculate differences of exon inclusion values (ΔΨs) across:

- For a matched cell type and region (HIPP and VIS), the developmental-time specific changes in exon inclusion were obtained by calculating pairwise ΔΨs. All comparisons that yielded ΔΨ ≥ 0.25 i.e., showed a 25% change in inclusion for an exon between two timepoints were reported
- For a given timepoint (P14, P21, and P28) and region (HIPP and VIS), the cell-type specific changes in exon inclusion were obtained by calculating pairwise ΔΨs. All comparisons that yielded ΔΨ ≥ 0.25 i.e., showed a 25% change in inclusion for an exon between two cell types were reported
- For a matched cell type at the adult timepont (P56), adult brain-region specific changes in exon inclusion were obtained by calculating pairwise ΔΨs. All comparisons that yielded ΔΨ ≥ 0.25 i.e., showed a 25% change in inclusion for an exon between two brain region were reported
- For a given brain region (HIPP, VIS, STRI, THAL, CEREB) at P56, the cell-type specific changes in exon inclusion were obtained by calculating pairwise ΔΨs. All comparisons that yielded ΔΨ ≥ 0.25 i.e., showed a 25% change in inclusion for an exon between two cell types were reported

The exons obtained from the four lineages above were classified as highly variable exons (hVEx). The ΔΨ for all pairwise comparisons of 44 triads, i.e., for 946 comparisons were then calculated for these hVEx were reported. To enable hierarchical clustering of this matrix of *comparisons x exons,* exons with too many NA values due to lack of depth for Ψ calculation in many triads were filtered out.

#### Identifying extremely variable exons (EVEx)

For the exons classified as highly variable (see above), and for each of the four lineages considered, we retained the comparison with the highest ΔΨ value. This yielded a matrix with 4 columns, one for each of the lineages, and 5931 rows, one for each hVEx. Of these, exons with ΔΨ≥ 0.75 in any of the four columns were retained. These were defined as extremely variable exons (EVEx), wherein at least one comparison between triads across the four lineages displayed 75% or more change in exon inclusion.

#### Obtaining developmental modalities of variability

Considering the second of the four lineages above, i.e., the cell-type specific differences in exon inclusion for a given timepoint and brain region, we calculated exon variability. For exons where we had sufficient depth to calculate the Ψ values for at least two cell types, we calculated the exon variability as the max(Ψ) – min(Ψ) for each timepoint. Thus,

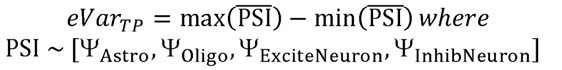

Then, we considered the first lineage from the hVEx paradigm above, i.e., developmental-time specific changes. Here, we looked at the change in variability between timepoint transitions, i.e., from P14 to P21, P21 to P28, and P28 to P56. If the change in eVar (Δ_eVar_) was less than 0.1 in all three transitions, indicating less than 10% change in cell-type specific variability over time, then the exon was classified as invariable (Fig S14).

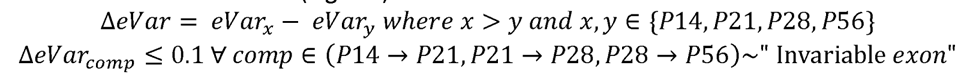

Otherwise, the exon was classified as variable, and the normalized matrix of *exon x eVar_TP_* was used for clustering and obtaining 9 developmental modalities.

#### Getting protein superfamily annotations per exon

We developed a computational pipeline to identify protein superfamily annotations per exon. For each exon, we used the genomeToProtein() function of the ensembldb package and extracted the ensembl ID, coordinates, and residue sequence of the protein identified. We filtered the obtained protein identifiers based on their corresponding ensembl transcript IDs and limited the search to the principal isoforms from APPRIS^7^, a database of curated principal and alternative isoforms which determines the principal isoforms based on the integrated analysis of the protein’s structure, function and cross-species conservation. For each protein sequence, we ran the SUPERFAMILY ^8, 9^ tool that utilizes Hidden Markov Model to identify the structural-defined SCOP protein domain families and the domain boundaries. The tool was implemented in InterProScan^10–12^. Protein regions not associated with domains were considered inter-domain linkers. Subsequently for each exon, the superfamily annotation associated with the protein residue for those coordinates was identified and used for further analysis.

#### Enrichment of protein superfamily annotations

We considered EVEx in clusters E1-E5 (Fig3), as well as background set consisting of exons with low variability across the entire dataset. For exons which had a domain associated with the coordinates of the coding sequence, we extracted the superfamily rather than individual and similar domains, given that the broader classification would allow for better grouping. We then counted the number of exons associated per superfamily and group, and reported a percentage. The superfamilies for which a value was obtained only for the background set were discarded, yielding a total of 55 superfamilies. For better interpretability, we retained superfamilies that were associated with at least 4% enrichment in any group, yielding a total of 15 superfamilies.

#### Mapping orthologous exons in human data

The TransMap ^13^ projection alignment algorithm was used to map exons between human and mouse assemblies. LASTZ^14^ genomics alignments between the human GRCh38 and mouse GRCm39 reference assemblies were used to map reference transcript annotations between assemblies. TransMap was used instead of UCSC Genome Browser liftOver^15^, as it produces base-level alignments, allowing observation of indels and other differences between the LASTZ chain and net alignments files^16^. These were obtained from the UCSC Genome Browser site, along with the below-mentioned programs to process them.

Syntenic genomic alignments were obtained by filtering the net files to obtain the syntenic nets using “netFilter −syn” and then using “netChainSubset -wholeChains” to obtain a set of syntenic chain alignments for mappings. GENCODE^17^ human v35 and mouse vM26 were mapped to the other assembly using the “pslMap” program.

#### Getting cell type variability in human hippocampus

We obtained Ψ values for each cell type in the human data using the same strategy as in for mouse data. Orthologous exons that were unambiguously and reciprocally mapped between human and mouse, shared sequence homology and length were selected. For EVEx in groups E1-E5 identified in mouse, exon Ψ values were obtained per cell type for exons where sufficient depth for the exons ortholog was available. Since only one brain region (HIPP) and timepoint (adult) were available, variability was defined as the max(Ψ) – min (Ψ) among the major cell types. Similarly, for all exons in the human data that had orthologs in mouse, Ψ values were obtained per major cell type and exons were classified as “highly” variable if the eVar ≥ 0.5 and invariable but alternative if eVar ≤ 0.2. To allow for a higher number of exons to be queried, we then obtained the Ψ values for the broad categories of neurons and glia, from both mouse and human data, and reported them.

#### Pseudotime trajectory analysis

Isoquant v3.1 was run on the ONT data and full-length isoforms were grouped by barcode to obtain a *isoform x cell* sparse matrix similar to the cellranger pipeline across the dataset. Cell barcodes corresponding to astrocytes and oligodendrocytes from the HIPP and VIS developmental lineage were isolated and a subset of a matrix containing these cellular barcodes as columns was obtained. This matrix was then processed using Seurat (v3.2.3) to obtain a UMAP representation of the cells. Each cell was colored according to the original short-read cell type assignments. Slingshot (v1.6)^18^ was then used to obtain a pseudotime trajectory along these clusters, with the initial point specified at OPCs. Lineages obtained were then reported.

#### Gene ontology analysis for genes associated with variable exon categories

For exons in the four hVEx categories (H1-H4), genes to which these exons belonged were extracted per category. Only unique genes per category were retained, meaning that if two exons from a gene belonged to two different categories, the gene was discarded from the analysis. Gene ontology (GO) biological process (BP) enrichment analysis was performed using the function enrichGO() from the clusterProfiler^19^ package. GO terms with qvalues ≤ 0.1 were reported, and the enrichment value was defined as the ratio of genes in the category being considered to those in the background. Similarly, for the invariable exons and the 9 variable developmental categories (G0-G9), unique genes containing the exons in each category were identified. GO-BP analysis was performed as above, using a list of brain-expressed genes obtained from SynGO^20^ as the background set. GO terms with qvalues ≤ 0.1 were reported, and the enrichment value was defined as the ratio of genes in the category being considered to those in the background. In the same vein, genes of adult brain-region specific EVEx (group E4) were identified and the same steps were performed.

#### Testing for exon coordination

Testing for exon coordination can be done at the pseudo-bulk level, or at the cell-type level. For every exon pair passing the criteria for sufficient depth, a 2 x 2 matrix of association for a given sample i.e., cell type or pseudo-bulk was generated. This matrix contained counts for inclusion of both exons (in-in), inclusion of the first exon and exclusion of the second (in-out), exclusion of the first exon and inclusion of the second (out-in), and exclusion of both exons (out-out).

The co-inclusion score of an exon was defined as the double inclusion (in-in) divided by the total counts for that exon pair. An exon pair was deemed “coordinated” was assessed using the *X^2^* test of association. The effect size was calculated as the |log10(oddsRatio)|. The odds-ratio was calculated by setting 0 values to 0.5, and dividing the product of double inclusion and double exclusion by the product of single-inclusion i.e. [(in-in) * (out-out)] / [(in-out) * (out-in)].

## Data availability

The summary of all data used for this study is/ will be made available on the Knowledge Brain Map (https://knowledge.brain-map.org/data/Z0GBA7V12N4J4NNSUHA/summary) which contains links to raw and processed data hosted on the Neuroscience Multi-Omic data archive (NeMO). All data supporting the findings of this study are provided within the paper and its supplementary information. Source data for the main figures can be found at https://github.com/noush-joglekar/biccn_tilgner_scisorseq/tree/main/data

## Code availability

The source code generated for this paper will be made publicly available at https://github.com/noush-joglekar/biccn_tilgner_scisorseq

