## Supplementary Materials for "Single-cell long-read mRNA isoform regulation is pervasive across mammalian brain regions, cell types, and development"

| **Timepoint** | **Brain region** | **Replicate** | **Sample ID** | **Total cells** | **Cells after QC** | **Median genes per cell** | **Median UMIs per cell** |
|---|---|---|---|---|---|---|---|
| P14 | HIPP | Rep1 | M5 | 14350 | 10046 | 2696 | 6042 |
| | | Rep2 | M6 | 10180 | 6711 | 3120 | 7651 |
| | VIS | Rep1 | M5 | 13661 | 10191 | 1865 | 3415 |
| | | Rep2 | M6 | 23317 | 19424 | 1673 | 3114 |
| P21 | HIPP | Rep1 | M1 | 11695 | 9355 | 1530 | 2803 |
| | | Rep2 | M5 | 13920 | 8271 | 2035 | 4560 |
| | VIS | Rep1 | M1 | 14231 | 10485 | 1272 | 2221 |
| | | Rep2 | M5 | 8655 | 8253 | 2251 | 4849 |
| P28 | HIPP | Rep1 | M1 | 13392 | 11067 | 2189 | 4746 |
| | | Rep2 | M2 | 11882 | 10240 | 2077 | 4224.5 |
| | VIS | Rep1 | M1 | 15320 | 12553 | 2144 | 4271 |
| | | Rep2 | M2 | 10046 | 8604 | 2504 | 5322 |
| P56 | HIPP | Rep1 | M1 | 6906 | 6494 | 1919 | 4354 |
| | | Rep2 | M2 | 12724 | 12124 | 1487 | 2839 |
| | VIS | Rep1 | M8 | 4733 | 4009 | 3285 | 8327 |
| | | Rep2 | M9 | 9564 | 7676 | 2196.5 | 5001 |
| | STRI | Rep1 | M1 | 8206 | 7270 | 1767 | 3739 |
| | | Rep2 | M2 | 7550 | 6618 | 2332 | 5073 |
| | THAL | Rep1 | M5 | 16145 | 13502 | 2116 | 4998 |
| | | Rep2 | M6 | 18278 | 16474 | 1985 | 4415 |
| | CEREB | Rep1 | M1 | 3676 | 3339 | 2971 | 9210 |
| | | Rep2 | M2 | 3870 | 3340 | 1993 | 3687 |

**Supplementary Table 1:** Sample details and quality control statistics for 10x single-cell short read sequencing libraries

| Timepoint | Brain region | Replicate | Sample ID | Total Reads | PolyA | Read length | Mapped and barcoded | CSMM |
|---|---|---|---|---|---|---|---|---|
| P14 | HIPP | Rep1 | M5 | 168303300 | 77.98 | 851.84 | 29959564 | 66.63 |
| | | Rep2 | M6 | 73356979 | 79.64 | 951.89 | 12957198 | 65.44 |
| | VIS | Rep1 | M5 | 53348838 | 77.05 | 842.75 | 9794529 | 67.11 |
| | | Rep2 | M6 | 104565321 | 80.78 | 823.28 | 12957198 | 65.44 |
| P21 | HIPP | Rep1 | M1 | 97967471 | 73.42 | 857.60 | 22819033 | 66.42 |
| | | Rep2 | M5 | 103148800 | 72.53 | 860.36 | 11736363 | 62.25 |
| | VIS | Rep1 | M1 | 68513664 | 73.67 | 860.39 | 17593692 | 66.32 |
| | | Rep2 | M5 | 50020807 | 71.21 | 876.01 | 6831543 | 64.54 |
| P28 | HIPP | Rep1 | M1 | 83183369 | 80.02 | 959.37 | 16422428 | 64.60 |
| | | Rep2 | M2 | 56387308 | 87.41 | 889.43 | 15610708 | 59.57 |
| | VIS | Rep1 | M1 | 82152770 | 78.77 | 918.15 | 17636898 | 64.80 |
| | | Rep2 | M2 | 85041433 | 86.52 | 908.46 | 21860630 | 60.53 |
| P56 | HIPP | Rep1 | M1 | 55104959 | 84.5 | 946.97 | 16908061 | 70.74 |
| | | Rep2 | M2 | 53430301 | 72.34 | 942.99 | 15552654 | 71.56 |
| | VIS | Rep1 | M8 | 76083018 | 81.31 | 968.55 | 21209929 | 74.06 |
| | | Rep2 | M9 | 157171800 | 84.73 | 1063.09 | 52197443 | 76.58 |
| | STRI | Rep1 | M1 | 72719845 | 78.02 | 965.52 | 21342316 | 73.25 |
| | | Rep2 | M2 | 58786362 | 74.27 | 953.77 | 16320143 | 71.25 |
| | THAL | Rep1 | M5 | 60842158 | 79.22 | 840.97 | 12024111 | 69.72 |
| | | Rep2 | M6 | 77752175 | 77.68 | 870.98 | 14836416 | 69.43 |
| | CEREB | Rep1 | M1 | 42192772 | 81.94 | 962.18 | 10509899 | 74.01 |
| | | Rep2 | M2 | 119513025 | 77.06 | 1014.49 | 30659173 | 65.77 |

**Supplementary Table 2:** Sample details and quality control statistics for unfragmented single-cell long-read libraries sequenced on Oxford Nanopore (ONT) PromethION

| Timepoint | Brain region | Replicate | Sample ID | Mean reads / SMRT cell | PolyA detected | Barcoded | Mean Length | CSMM Reads |
|---|---|---|---|---|---|---|---|---|
| P14 | HIPP | Rep1 | M5 | 2816040 | 88.73 | 46.56 | 898.67 | 67.11 |
| | | Rep2 | M6 | 3921610 | 87.07 | 36.88 | 1002.27 | 66.98 |
| | VIS | Rep1 | M5 | 2599560 | 83.14 | 38.23 | 929.94 | 70.59 |
| | | Rep2 | M6 | 3366700 | 91.43 | 49.33 | 887.56 | 68.78 |
| P28 | HIPP | Rep1 | M1 | 2698694 | 81.98 | 51.42 | 1004.28 | 70.12 |
| | | Rep2 | M2 | 3074310 | 88.25 | 55.96 | 992.39 | 67.03 |
| | VIS | Rep1 | M1 | 2820433 | 84.14 | 53.10 | 1059.71 | 69.07 |
| | | Rep2 | M2 | 1990540 | 86.64 | 52.27 | 1012.07 | 68.34 |
| P56 | HIPP | Rep1 | M1 | 2802050 | 91.03 | 70.28 | 1155.99 | 70.71 |
| | | Rep2 | M2 | 2620720 | 80.59 | 69.51 | 1071.05 | 72.66 |
| | STRI | Rep1 | M1 | 2212459 | 87.66 | 70.75 | 1099.93 | 70.99 |
| | | Rep2 | M2 | 2083750 | 84.68 | 67.96 | 1103.04 | 72.75 |
| | CEREB | Rep1 | M1 | 2755640 | 89.90 | 59.06 | 1125.40 | 72.91 |
| | | Rep2 | M2 | 2705560 | 85.64 | 61.03 | 1137.35 | 65.39 |
| | THAL | Rep1 | M5 | 3724854 | 85.10 | 41.96 | 1012.66 | 72.56 |
| | | Rep2 | M6 | 1349837 | 85.26 | 45.48 | 1026.63 | 71.79 |

**Supplementary Table 3:** Sample details and quality control statistics for unfragmented single-cell long-read libraries sequenced on Pacific Biosciences (PacBio) HiFi Sequel II

| Timepoint | Brain region | Replicate | Sample ID | min Genes | max Genes | min UMIs | max UMIs | mito cutoff | dims | resolution |
|---|---|---|---|---|---|---|---|---|---|---|
| P14 | HIPP | Rep1 | M5 | 500 | 6000 | 1000 | 25000 | 20 | 30 | 0.6 |
| | | Rep2 | M6 | 500 | 6000 | 1000 | 30000 | 30 | 30 | 0.6 |
| | VIS | Rep1 | M5 | 500 | 6000 | 1000 | 20000 | 20 | 30 | 0.6 |
| | | Rep2 | M6 | 500 | 6000 | 1000 | 25000 | 30 | 30 | 0.6 |
| P21 | HIPP | Rep1 | M1 | 500 | 5000 | 1000 | 15000 | 25 | 30 | 0.6 |
| | | Rep2 | M5 | 500 | 6000 | 1000 | 25000 | 20 | 30 | 0.6 |
| | VIS | Rep1 | M1 | 500 | 6000 | 1000 | 18000 | 40 | 30 | 0.6 |
| | | Rep2 | M5 | 1000 | 6000 | 1000 | 30000 | 30 | 25 | 0.6 |
| P28 | HIPP | Rep1 | M1 | 500 | 6000 | 1000 | 25000 | 18 | 30 | 0.6 |
| | | Rep2 | M2 | 500 | 6000 | 1000 | 25000 | 15 | 30 | 0.6 |
| | VIS | Rep1 | M1 | 500 | 6000 | 1000 | 25000 | 18 | 30 | 0.6 |
| | | Rep2 | M2 | 500 | 7000 | 1000 | 30000 | 18 | 25 | 0.6 |
| P56 | HIPP | Rep1 | M1 | 500 | 7000 | 1000 | 20000 | 15 | 30 | 0.6 |
| | | Rep2 | M2 | 500 | 7000 | 1000 | 20000 | 15 | 30 | 0.6 |
| | VIS | Rep1 | M8 | 1000 | 7500 | 1000 | 40000 | 7 | 25 | 0.6 |
| | | Rep2 | M9 | 1000 | 7500 | 1000 | 40000 | 10 | 25 | 0.6 |
| | STRI | Rep1 | M1 | 500 | 5000 | 1000 | 20000 | 15 | 25 | 0.6 |
| | | Rep2 | M2 | 500 | 6000 | 1000 | 25000 | 18 | 25 | 0.6 |
| | THAL | Rep1 | M5 | 400 | 7500 | 1000 | 40000 | 35 | 30 | 0.6 |
| | | Rep2 | M6 | 400 | 7500 | 1000 | 40000 | 35 | 30 | 0.6 |
| | CEREB | Rep1 | M1 | 500 | 6000 | 1000 | 25000 | 20 | 20 | 0.5 |
| | | Rep2 | M2 | 500 | 6000 | 1000 | 25000 | 20 | 20 | 0.5 |

**Supplementary Table 3:** Parameters used in the processing of single-cell short read data

| Subject ID | Age | Hemisphere | Subject Sex | Race | PMI (hours) | RIN | Clinical Brain Diagnosis | Sample ID assigned |
|---|---|---|---|---|---|---|---|---|
| 1364 | 33 | Left | Female | Black or African-American | 20 | 99.99 | Unaffected Control | f2 |
| 1114 | 31 | Left | Male | White | 15 | 7 | Unaffected Control | m2 |
| 1539 | 33 | Left | Female | White | 23 | 7.1 | Unaffected Control | f1 |
| 5398 | 36 | Left | Male | White | 24 | 8.3 | Unaffected Control | m3 |
| 4676 | 40 | Left | Female | Black or African-American | 15 | 7.1 | Unaffected Control | f3 |
| 6096 | 28 | Left | Male | White | 7 | 8.3 | Unaffected Control | m1 |

**Supplementary Table 5:** Sample IDs and subject details for single-nuclei hippocampal samples

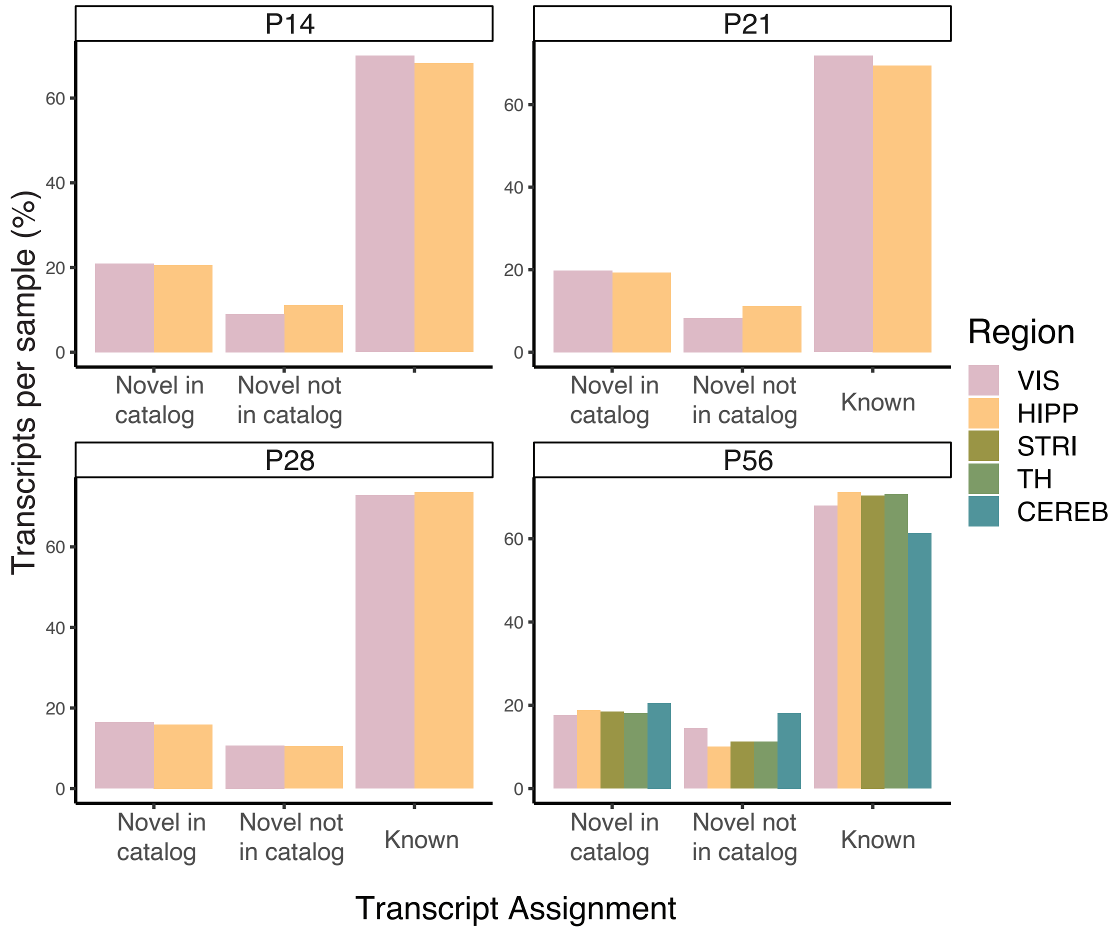


**Supplementary Figure 1: Transcript assignments for ONT data.** Barplots of the percentage of transcript in each sample, averaged over replicates, and classified by IsoQuant as novel in catalog, novel not in catalog or Known. Color of bar indicates region of origin. Each sub-panel represents the developmental timepoint from which the samples were collected.

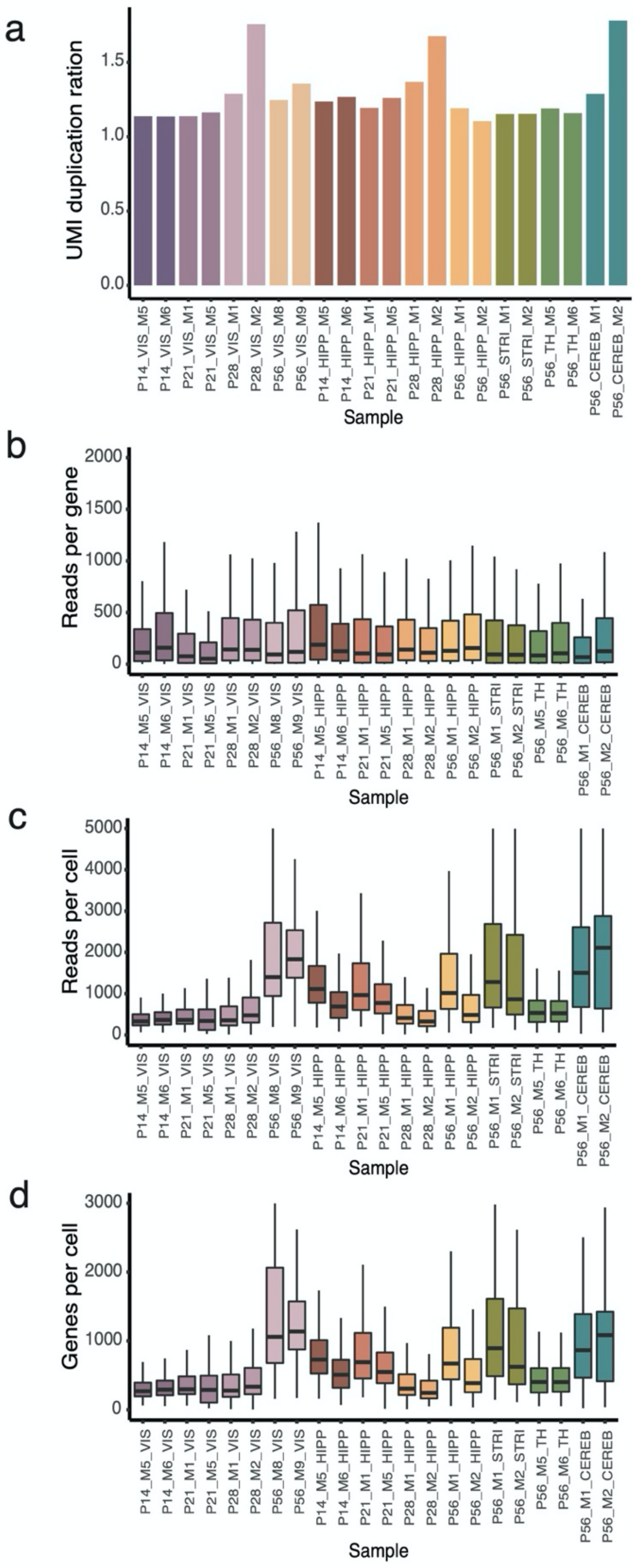


**Supplementary Fig 2: Quality control for ONT long reads. (a)** Barplot of the UMI duplication ratio per sample indicated on the x-axis. **(b)** Boxplot of the number of spliced, barcoded, umi deduplicated reads per gene in each sample. **(c)** Same as (b) but with reads per cell in each sample. **(d)** Same as (b) but with number of genes per cell in each sample. Color of bar represents sample of origin, each replicate has the same color.

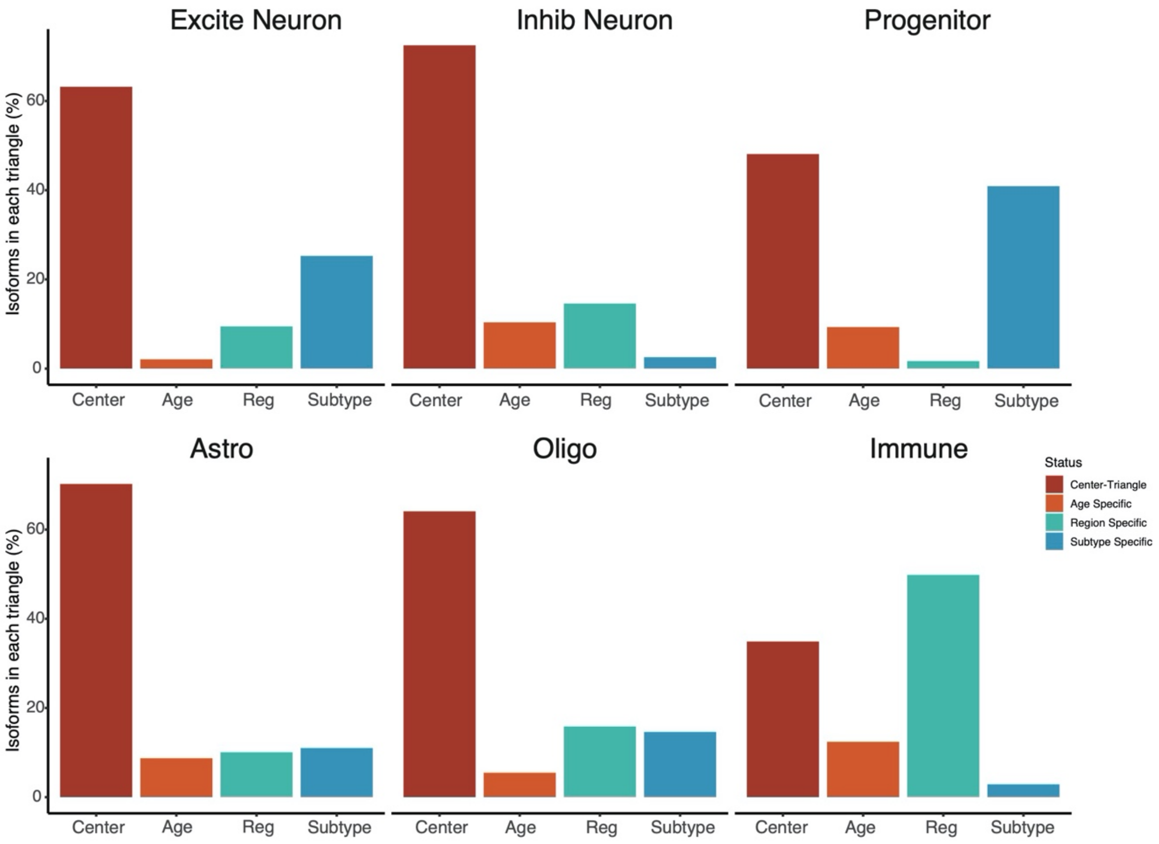


**Supplementary Fig 3: Variability assignment for full-length isoforms.** Barplot of the percent of isoforms per cell type that was assigned to each triangle in the cell-type specific ternary plot. Red: Center triangle, orange: age-specific isoforms, teal: region specific isoforms, blue: Cell subtype specific isoforms. Each panel represents cell type in which isoform variability was assessed.

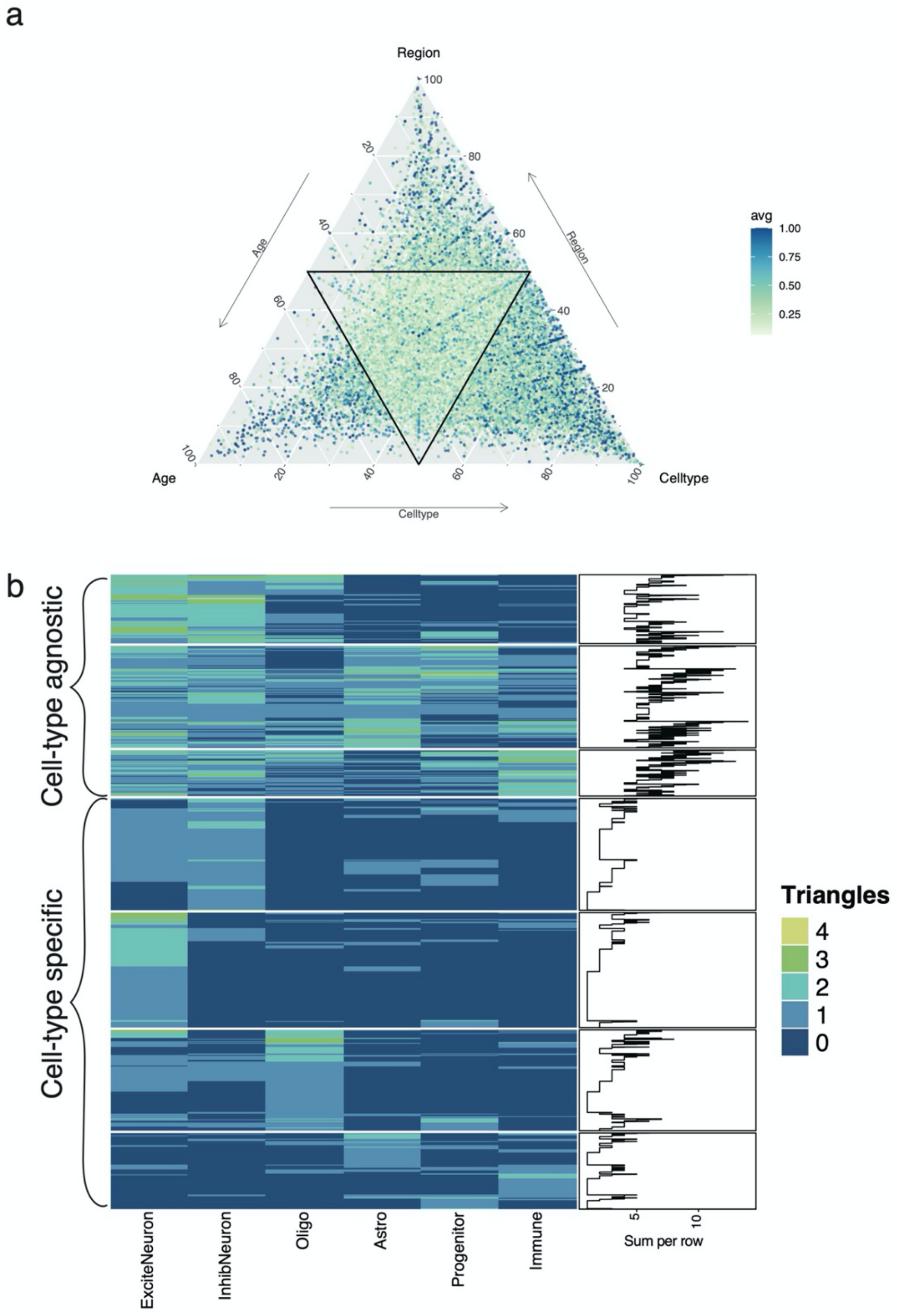


**Supplementary Fig 4: Isoform variability in pseudobulk and hypervariable genes (a)** Ternary plot of the isoform variability in pseudobulk with triangles representing age, region, or cell type specificity per isoform at each vertex. Each point is a single isoform, and color of point represents the average raw variability value along all three axes. **(b)** Heatmap of hypervariable genes per cell type. Color of each tile in the heatmap indicates the number of triangles in which an isoform was found to be highly variable (variability $\geq$ 0.25) for a given cell type. Annotation bar on the right denotes the line plot of the sum of the number of triangles per row. High values indicate genes wherein isoforms are regulated along multiple axis along multiple cell types and are therefore highly variable and cell-type agnostic. Low values / non-zero values in a few cell-types indicates highly specialized isoforms with cell-type specificity.

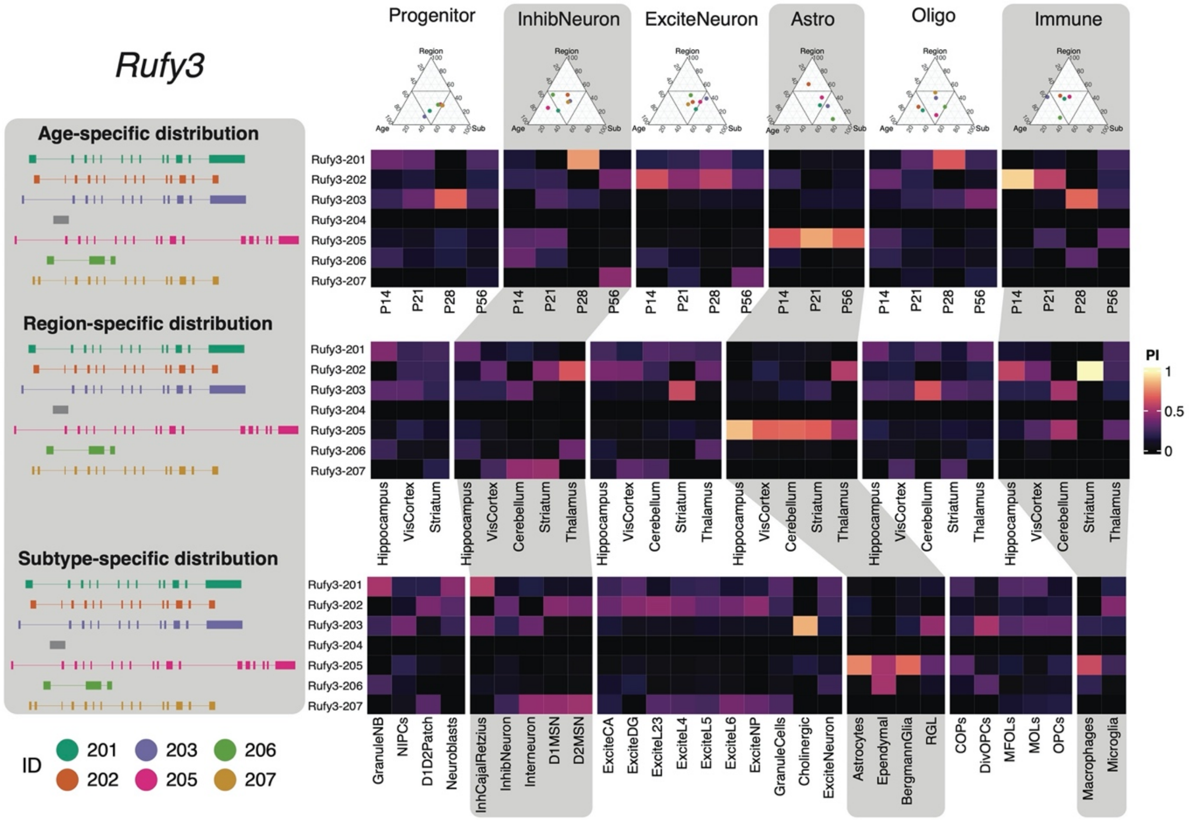


**Supplementary Fig 5: Isoform regulation in *Rufy3.*** Left panel in grey showing seven annotated isoforms of the gene *Rufy3* with three sections indicating age, region, and subtype specific distributions to be represented in the heatmaps on the right. Ternany plots on the right are shown per cell type, with the isoform variability value per isoform represented in the plot, where each point represents a *Rufy3* isoform colored according to the legend on the bottom left. Each heatmap shows an isoform (rows) per cell-type in each developmental stage as columns (top section), each brain region as columns (middle section) and each cell subtype as columns (bottom section). Color of tile shows the percent inclusion (Π).

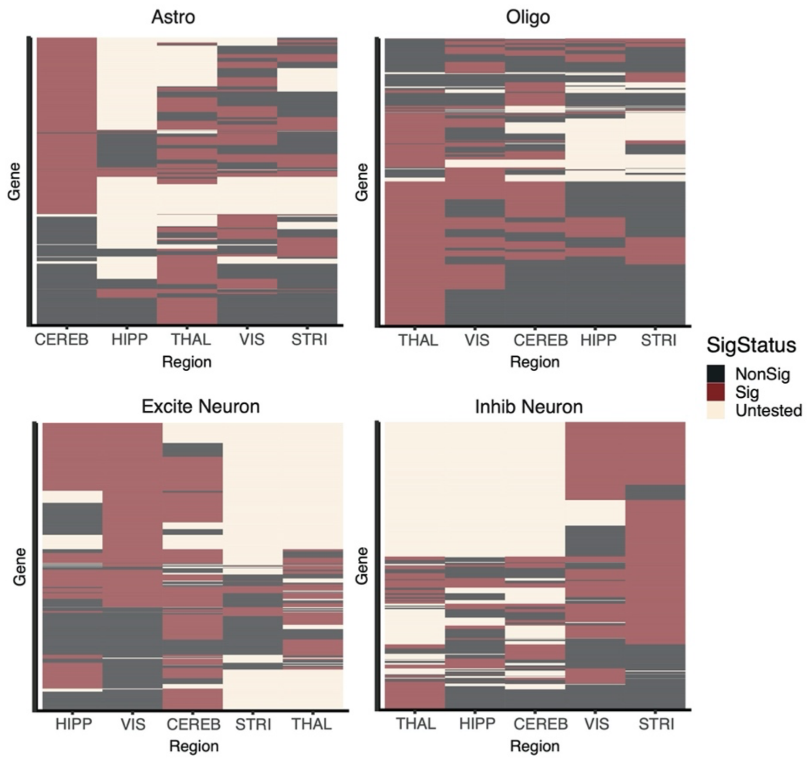


**Supplementary Fig 6: Differential isoform expression results summary per cell type.** Heatmap summarizing the results of differential isoform expression tests of one brain region (represented as columns) versus an aggregate of all others for each major cell type at P56. Color of tile indicates whether the gene was considered significant i.e., FDR $\leq 0.05$ and $\Delta\Pi \geq 0.1$ (maroon), non-significant (black), or was not tested (tan).

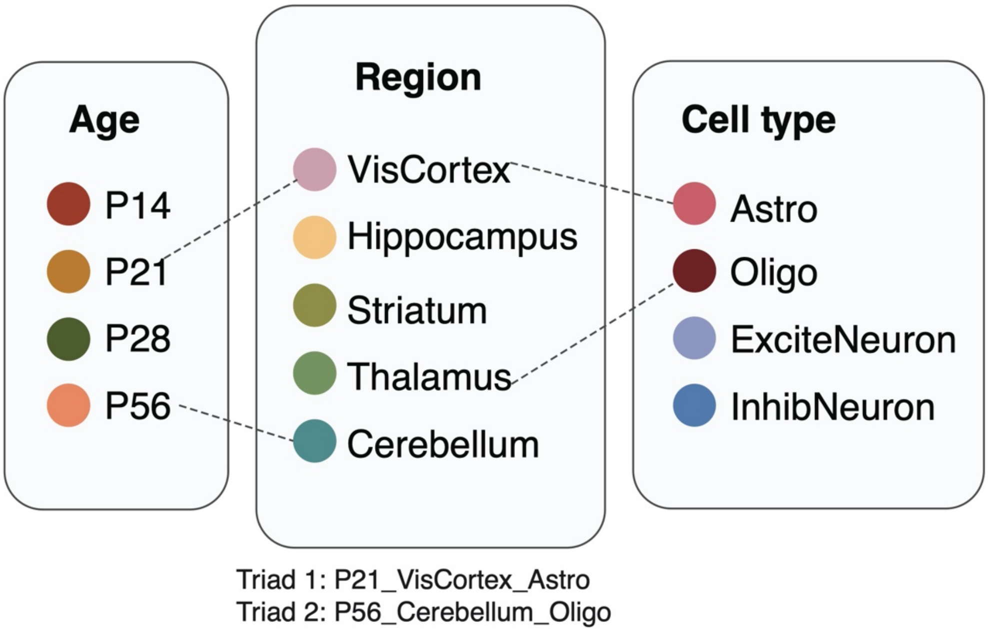


**Supplementary Fig 7: Definition of a triad.** A triad can be defined by the age, region, and major cell type of origin, with counts being averaged over replicates. This results in 44 triads given sample constraints.

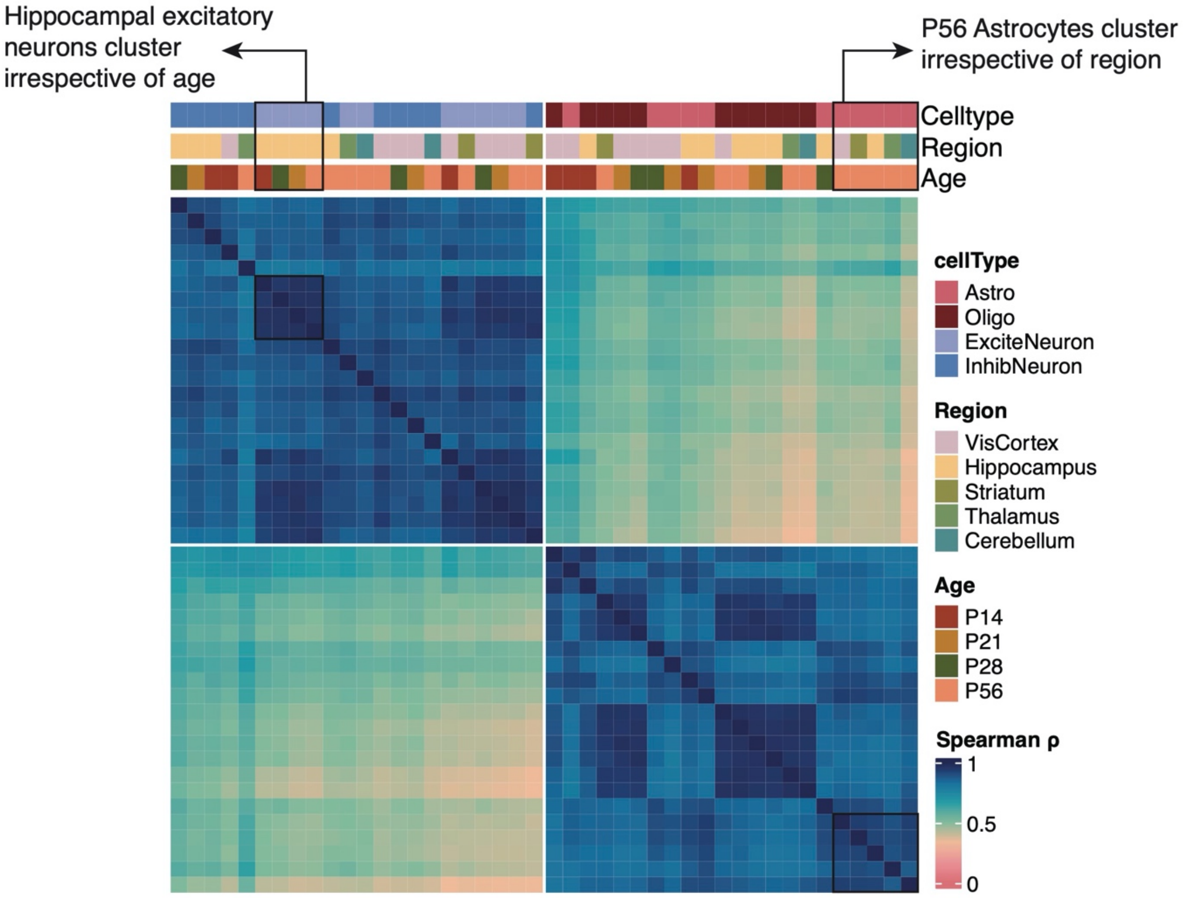


**Supplementary Fig 8: Correlation of triad PSIs.** Heatmap representing the spearman correlation values of triad exon $\Psi$ values across all measurable exons. Color of tile represents the spearman $\rho$ Annotation bars represent the major cell type (top), brain region (middle), and age (bottom) contributing to each of the 44 triads. Black outlines indicate astrocytes at P56 which have clustered together regardless of brain region, and excitatory neurons in the hippocampus which have clustered together regardless of timepoint.

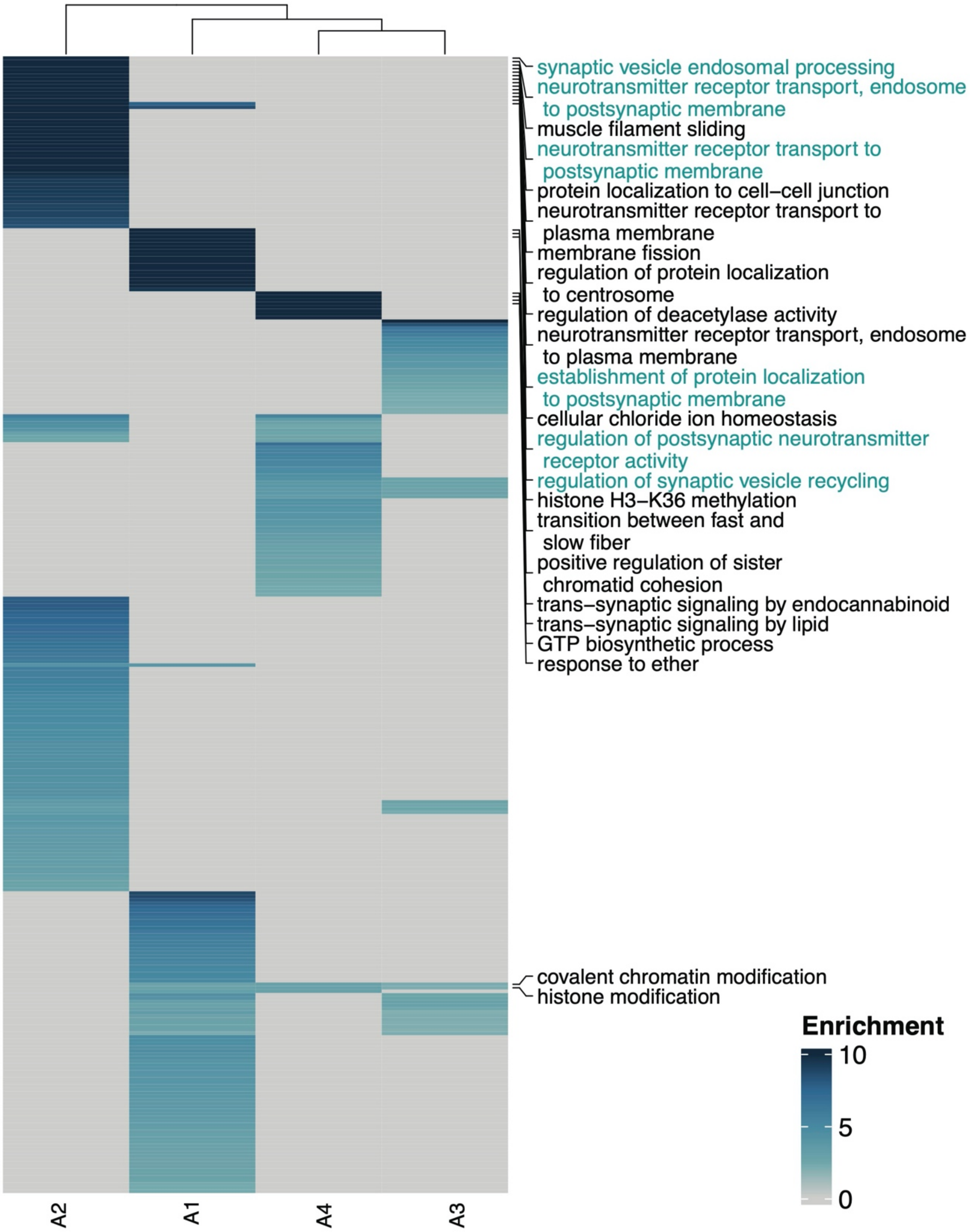


**Supplementary Fig 9: Gene ontology of hVEx groups.** Heatmap of the biological process gene ontology (GO-BP) enrichment terms for each of the four categories of highly variable exons (hVEx). Color of tile indicates the level of enrichment in categories compared to the background. GO terms with very high enrichment observed in a single category, or medium enrichment observed across multiple categories are reported.

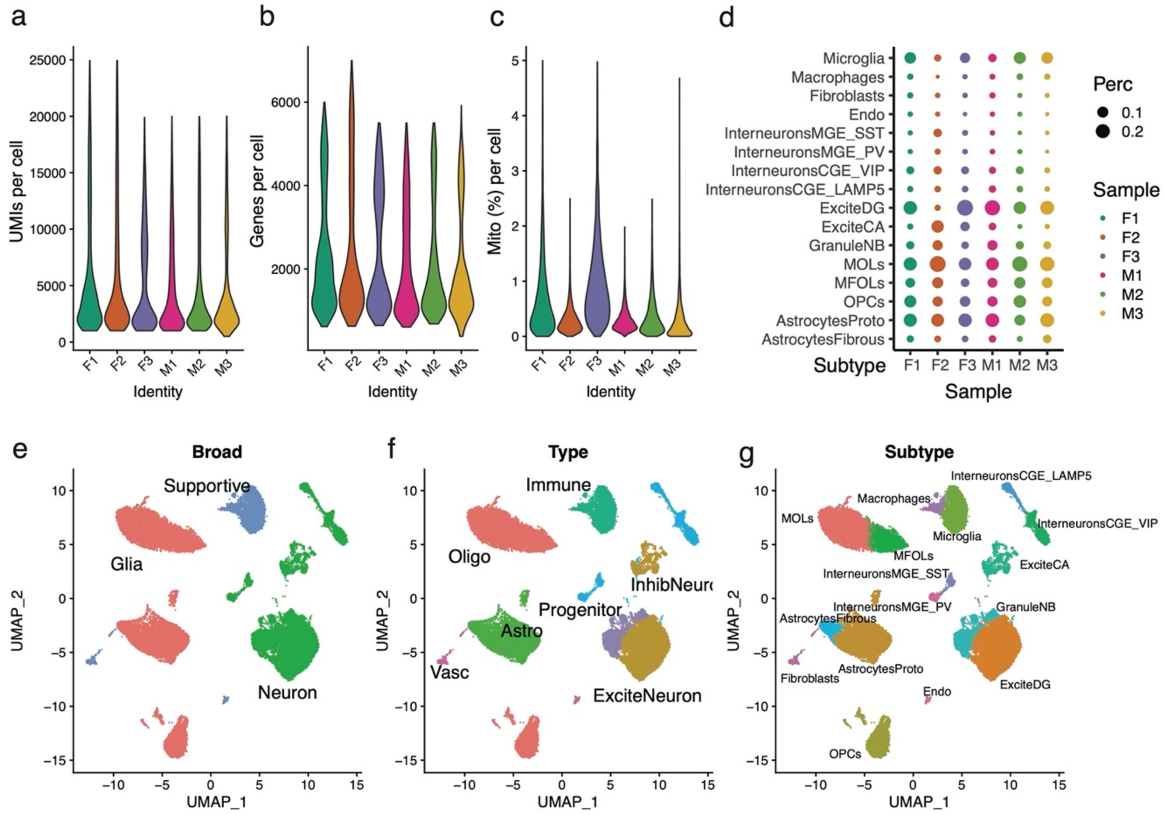


**Supplementary Fig 10: Processing of short-read snRNA human hippocampal data. (a)** Violin plots of the UMIs per nucleus for each of the six hippocampal samples indicated on the x-axis. (b) Same as in (a) but showing a distribution of genes per cell nucleus (c) Same as in (a) but showing the distribution of the percentage of mitochrondrial reads sequenced per cell nucleus. (d) A dot plot indicating the percentage of nuclei per sample (x-axis) belonging to a cell subtype (y-axis). Size of dot indicates percentage while color of dot indicates sample. (e) UMAP embedding of all nuclei from the six samples clustered together after controlling for batch effects Each point represents a single cell nucleus and color of cluster denotes the broad cell type i.e., green: neurons, red: glia, and blue: supportive cells. (f) Same UMAP as in (e) but colored by cell type. (g) Same UMAP as in (e) but colored by cell subtype.

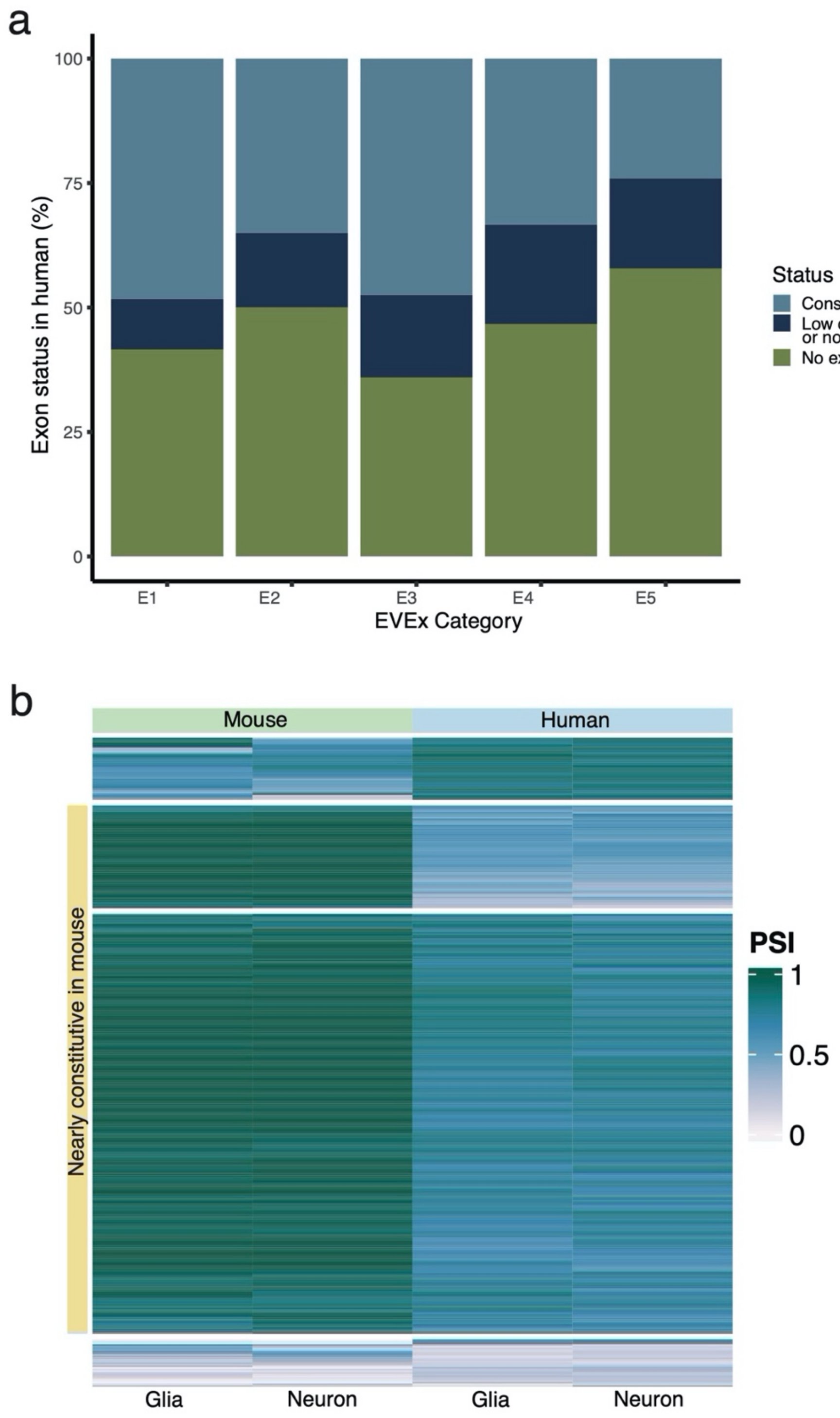


**Supplementary Fig 11: Exon conservation between mouse and human (a)** Barplots for each group of extremely variable exons (EVEx, E1- E5) indicated on the x-axis with the percentage of exons on the y-axis. Bars are split according to whether the exon has an ortholog and has a calculable $\Psi$ in human data (light blue), does not have an ortholog in human mapped with high-confidence (navy), or has an ortholog but is too lowly expressed to calculate a $\Psi$ value. **(b)** Heatmap of $\Psi$ values in glia and neurons in mouse (left side) versus humans (right size) for exons that are lowly variable in humans. Yellow annotation bar on the left indicates exons in which $\Psi$ values are close to 1 in mouse, indicating that the exons are nearly constitutively included in mouse.

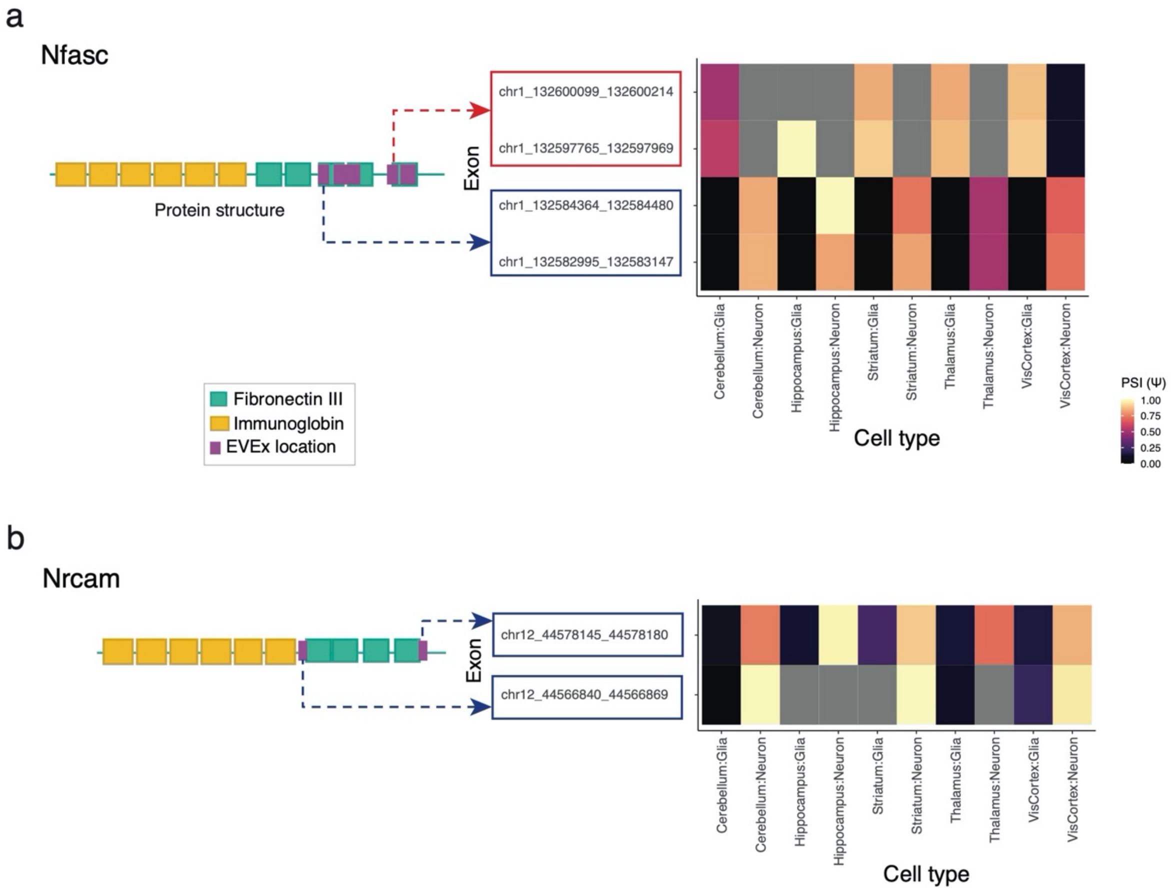


**Supplementary Fig 12: Domain architecture and exon inclusion values for NFASC and NRCAM. (a)** *Left* Domain information for the NFASC protein with immunoglobin (IG)-like domains in yellow followed by Fibronectin Type III (Fn3) domains in teal. Highlighted parts (purple) denote EVEx with cell type specificity. *Right* Heatmap of exon inclusion (PSI) values for the four EVEx identified for the *Nfasc* gene. Top two exons show a clear preference for glial inclusion while bottom two are highly included in neurons for all five brain regions at P56. Grey tiles indicate NA values. **(b)** Same as in (a) but for the gene *Nrcam* where both exons are highly included in neurons.

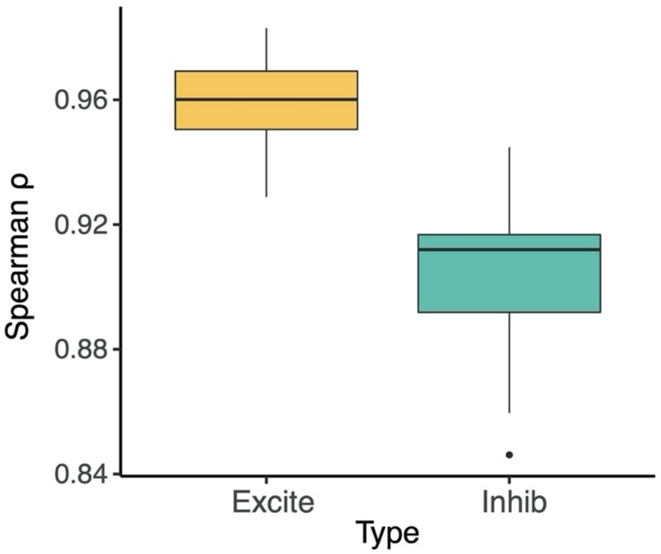


**Supplementary Fig 13: Correlation within excitatory and inhibitory neurons.** Boxplot of the pairwise spearman correlation (ρ) values of the exon inclusion values across all subtypes of excitatory neurons in VIS in all developmental timepoints (yellow), and same for inhibitory neurons (teal).

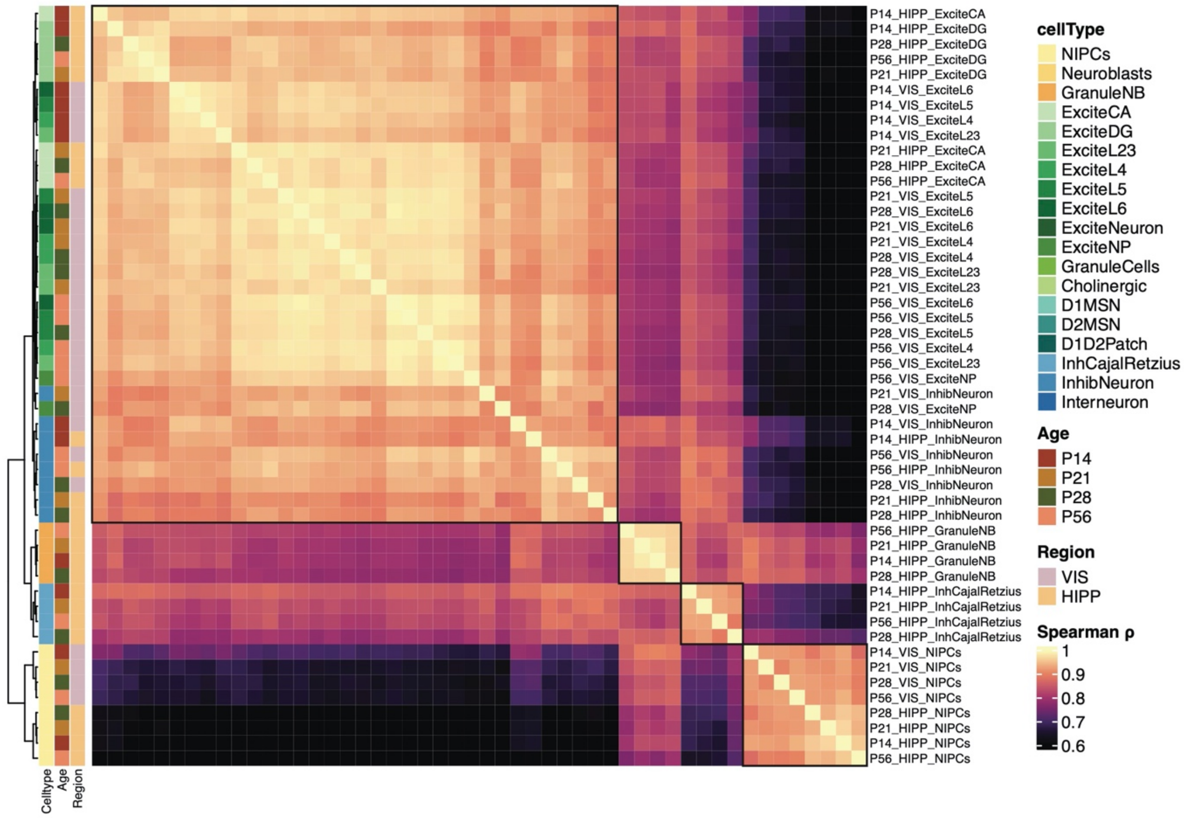


**Supplementary Fig 14: Correlation of neuronal subtype Ψ.** Heatmap of the pairwise spearman correlation (ρ) values of the exon Ψ across all subtypes of excitatory and inhibitory neurons in HIPP and VIS considered together. Annotation bars on left indicate the cell subtype (left-most), age (middle), and region (right-most) of origin for each subtype considered. Black boxes outline the major splits in the dendrogram.

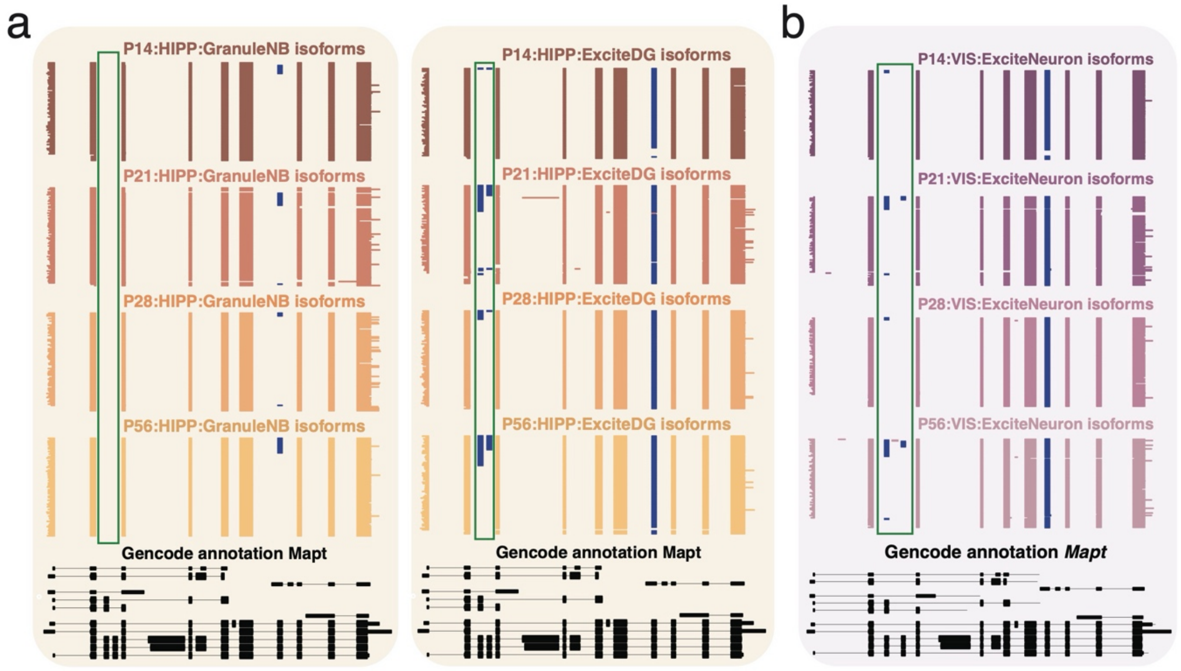


**Supplementary Fig 15: Developmental isoform expression of *Mapt* showing developmental isoform variability. (a)** ScisorWiz plot showing the isoforms for the gene *Mapt* for granule neuroblasts (left) and dentate gyrus excitatory neurons (right) in the hippocampus colored and split by age. Each line indicates a unique cDNA molecule, with clustered chunks denoting exons. Alternative exons denoted in blue. The first two alternative exons are outlined with a green box. Black lines on the bottom indicates GENCODE annotated transcripts. **(b)** Plot with a similar structure as in (a) but for visual cortex excitatory neurons.

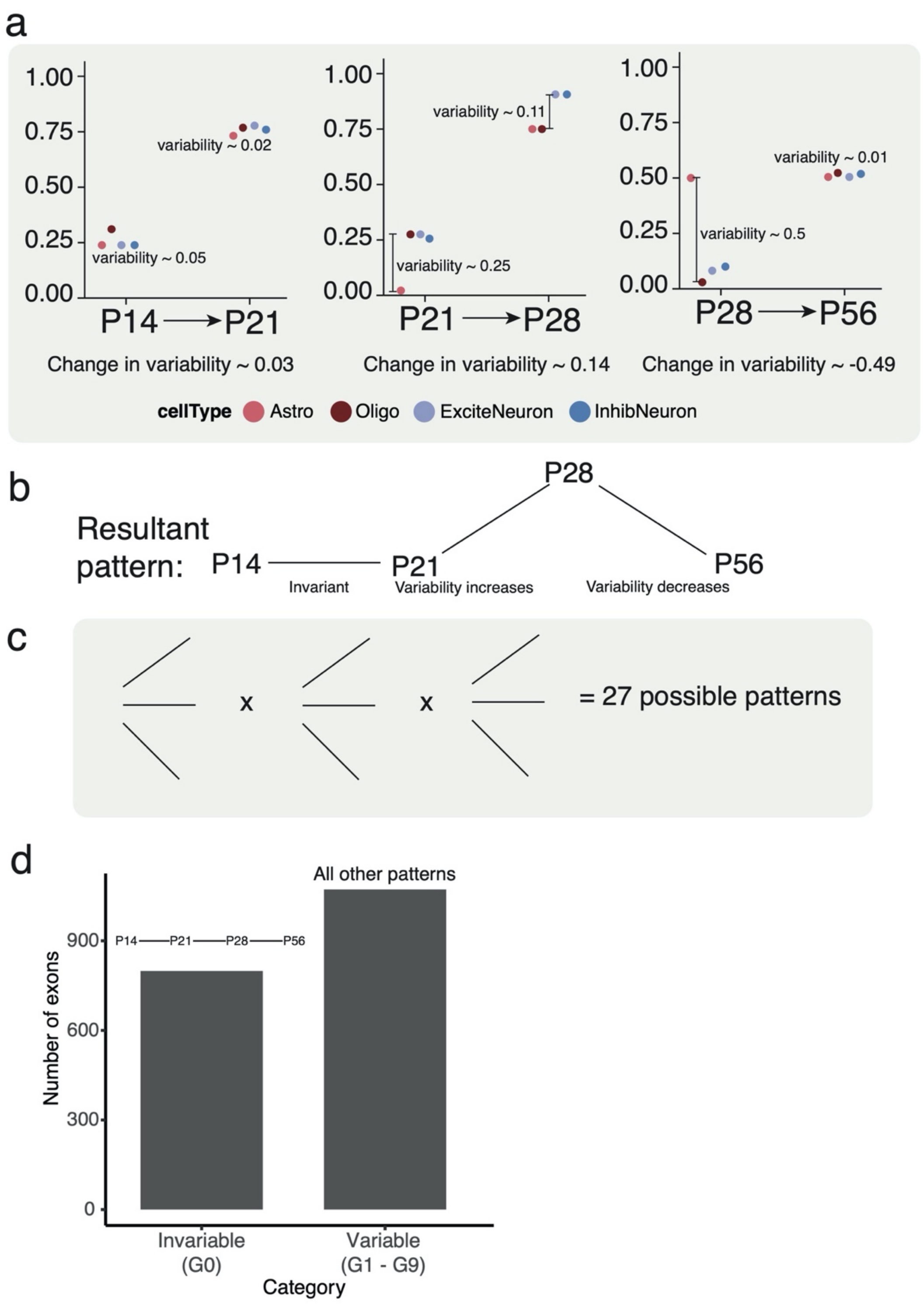


**Supplementary Fig 16: Defining patterns of developmental variability. (a)** Schematic representing the change in variability for major cell types in each of the three developmental transitions indicated on the x-axis. X-axis indicates time point and y-axis shows the $\Psi$ value. **(b)** The pattern of variability resulting from the hypothetical schematic shown in (a). **(c)** Given that the variability can go up, down, or remain fairly constant, and there are three developmental transitions under considerations, 27 patterns of variability are theoretically possible. **(d)** Under the definition that the variability $\leq 0.1$ for each of the three transitions, a barplot showing the number of exons defined as invariable versus variable. The variable exons are then used to obtain the 9 developmental patterns of variability.

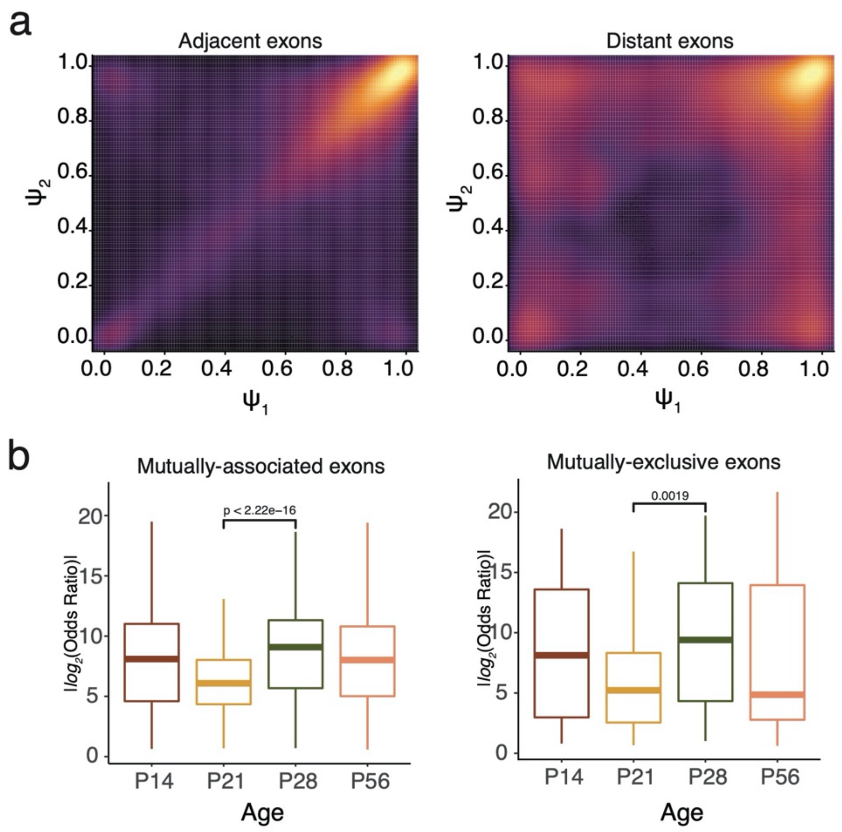


**Supplementary Fig 17: Inclusion patterns for pairs of exons. (a)** A smoothed scatterplot showing the individual Ψ of pairs of adjacent exons (*left*) which are often included or skipped together, and distant exons (*right*) which can be mutually exclusive. Black/ purple indicates low enrichments of points while yellow indicates high enrichment. **(e)** Boxplots of the effect size of coordination for pairs of exons. Y axis show the absolute value of the $\log_2$ odds ratio for mutually associated exons (*left*) and mutually exclusive exons (*right*) for each of the developmental timepoints indicated on the x-axis. Higher absolute odds ratio indicates tighter coordination. P-value obtained from two-sided Wilcoxon rank sum test

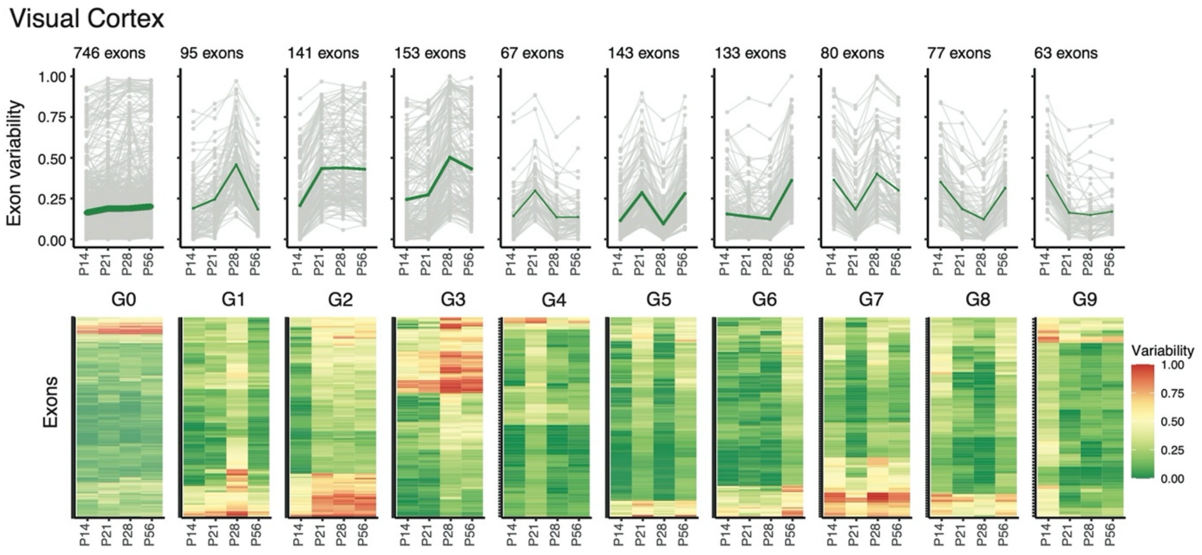


**Supplementary Fig 18: Patterns of developmental variability in the mouse visual cortex.** Line plots showing the variability value for exons in the invariable category (G0) and the nine patterns of variability identified (G1-G9) on the y-axis and the four timepoints on the x-axis. The number of exons in each category for visual cortex is indicated above. Green line connects the mean variability value per group and time point (*top*). Heatmap of the exon variability for each category (G0-G9) and each time point indicated on the x-axis (*bottom*)

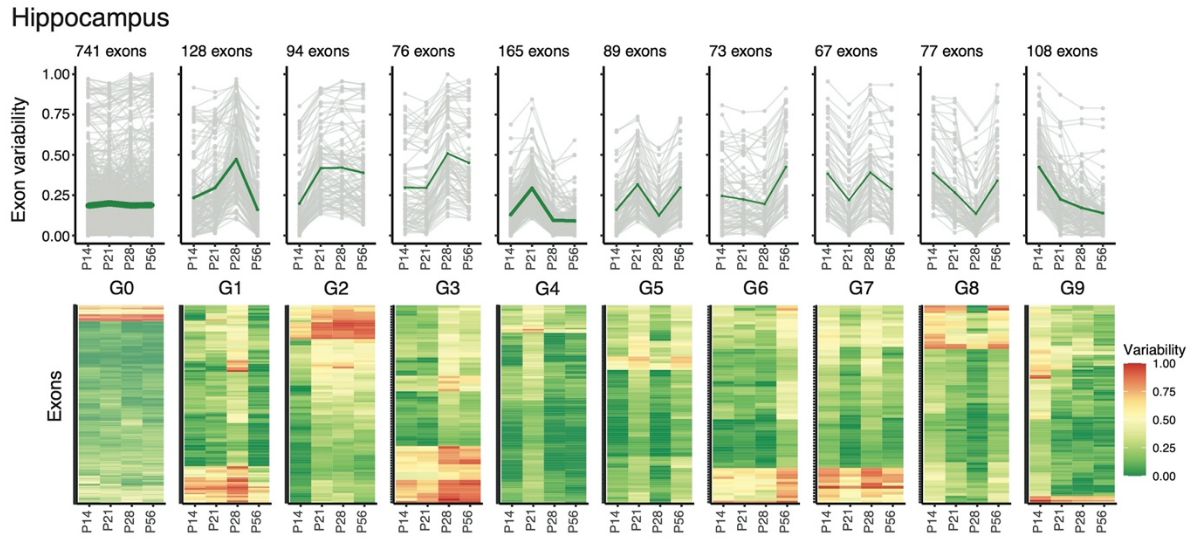


**Supplementary Fig 19: Patterns of developmental variability in the mouse hippocampus.** Line plots showing the variability value for exons in the invariable category (G0) and the nine patterns of variability identified (G1-G9) on the y-axis and the four timepoints on the x-axis. The number of exons in each category for hippocampus is indicated above. Green line connects the mean variability value per group and time point (*top*). Heatmap of the exon variability for each category (G0-G9) and each time point indicated on the x-axis (*bottom*)

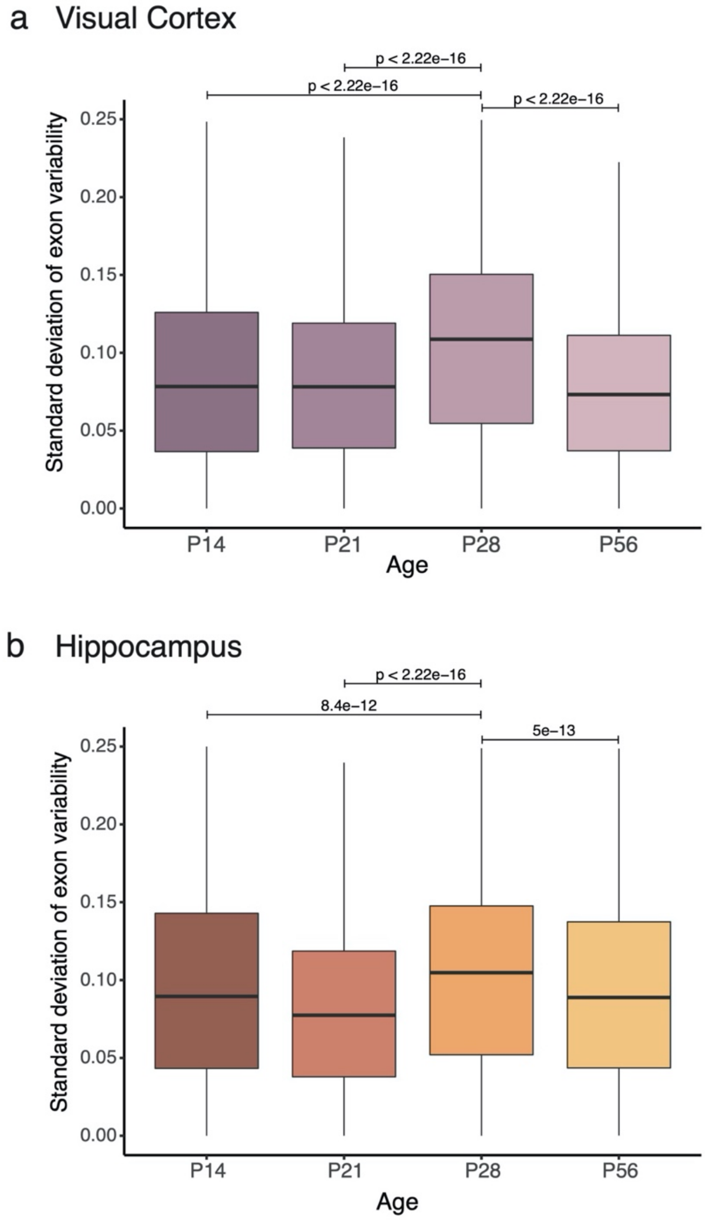


**Supplementary Fig 20: P28 as a critical timepoint for exon variability between cell types (a)** Boxplots showing the standard deviation of exon variability between the major cell types in the visual cortex on the y-axis, at each timepoint indicated on the x-axis. **(b)** Same as in (a) but for hippocampus.

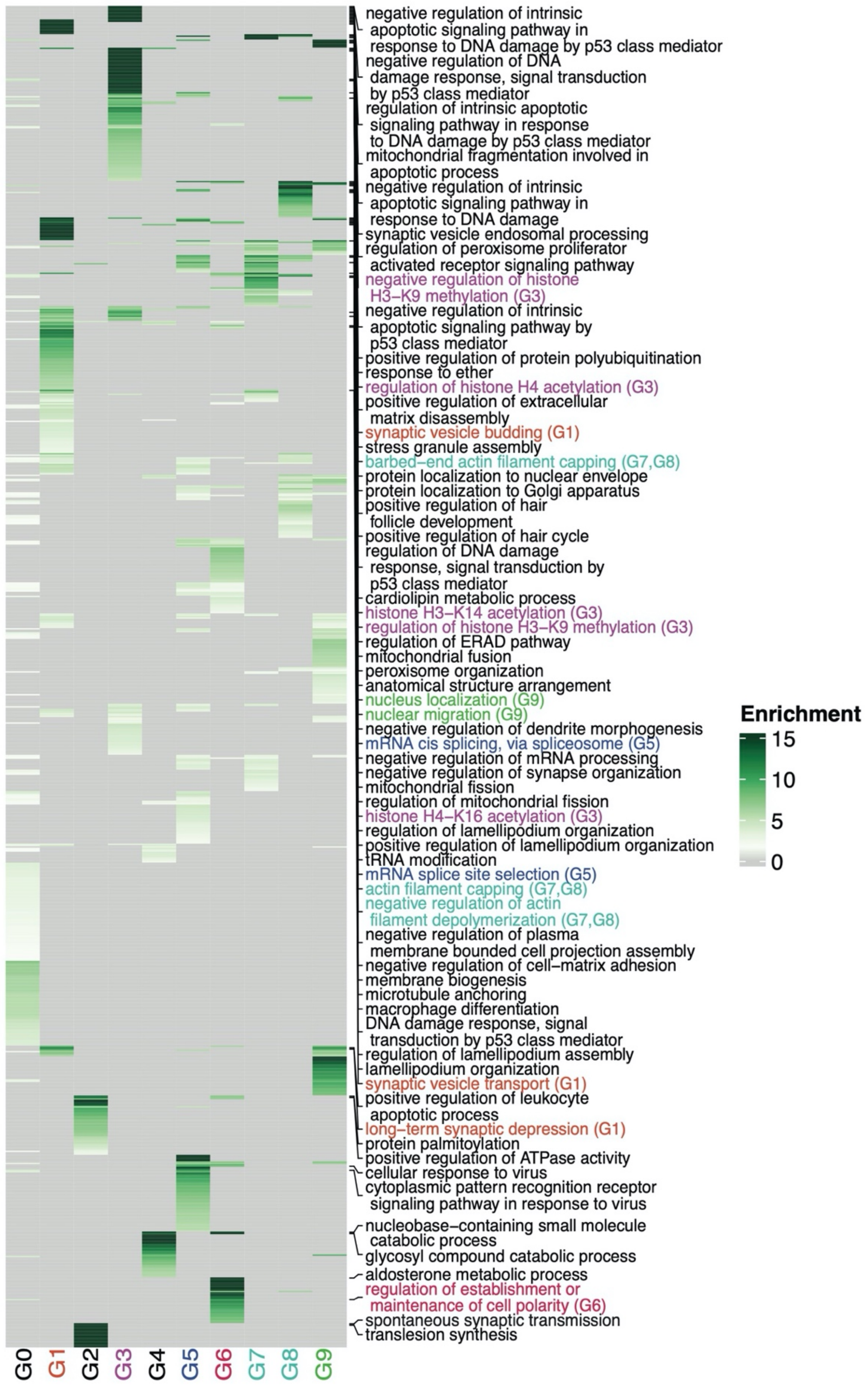


**Supplementary Fig 21: Gene ontology enrichment for the developmental patterns of variability.** Heatmap of the biological process gene ontology (GO-BP) enrichment terms for each of the nine variable and one invariable (G0) patterns of developmental variability of exon inclusion. Color of tile indicates the level of enrichment in categories compared to the background. GO terms with very high enrichment observed in a single category, or medium enrichment observed across multiple categories are reported. Some terms and the group they belong to are highlighted in colors to aid the reader.

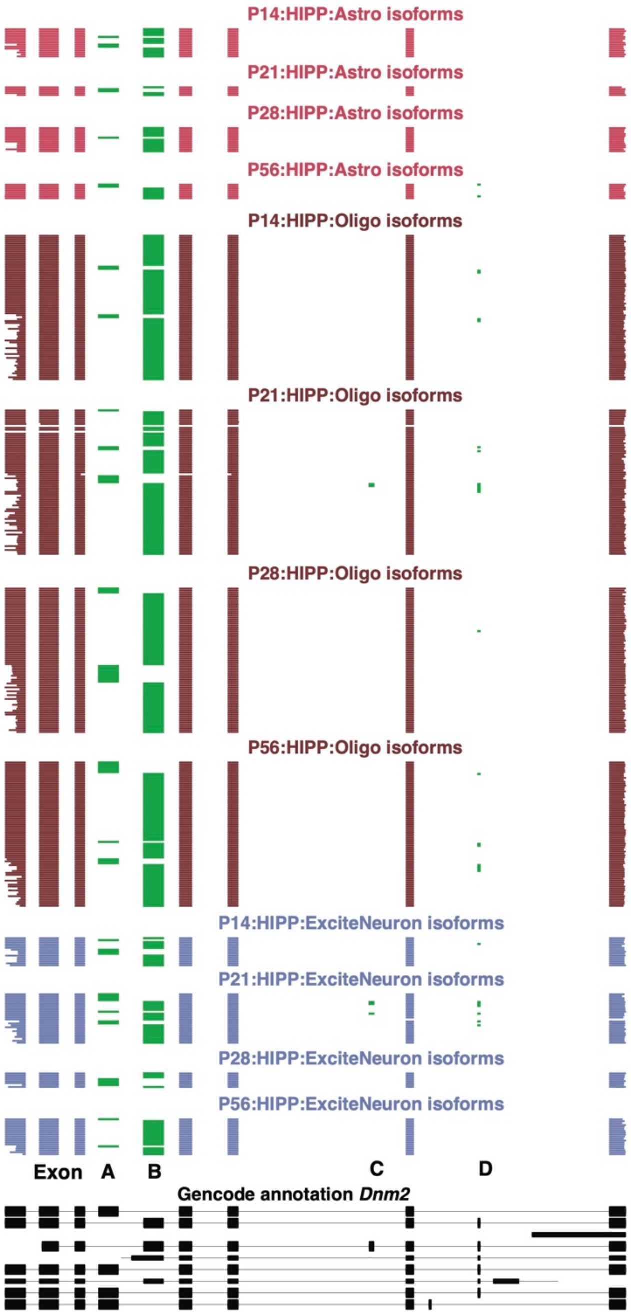


**Supplementary Fig 22: Isoform expression of *Dnm2* in hippocampus.** ScisorWiz plot showing the isoforms for the gene *Dnm2* for astrocytes (top, pink), separated by timepoint, oligodendrocytes (middle, maroon), and excitatory neurons (bottom, blue). Each line indicates a unique cDNA molecule, with clustered chunks denoting exons. Alternative exons are colored in green and are represented on the bottom. Black lines on the bottom indicates GENCODE annotated transcripts. Compare to Fig 6d-e
